# Lung adventitial fibroblasts support type 2 Tregs to shape the response to influenza infection

**DOI:** 10.64898/2026.04.07.716742

**Authors:** Anthony A. Chang, Joanna Balcerek, Sofia E. Caryotakis, Kennedi B. Pyper, Peri Matatia, Elina B. Wells, Tatsuya Tsukui, Nathan A. Ewing-Crystal, Eric Dean Merrill, Nick M. Mroz, Madelene W. Dahlgren, Kelly M. Cautivo, Marcela T. Taruselli, Hiromi Nakao-Inoue, Anna V. Molofsky, Dean Sheppard, Ari B. Molofsky

## Abstract

Viral pneumonias are lung infections that lead to both short- and long-term complications, including acute respiratory distress syndrome (ARDS) and pulmonary fibrosis. Lung stromal–immune interactions balance pathogen clearance with excessive immune-mediated injury, and healthy lung repair with necessary scarring. Here, we define the lung spatiotemporal stromal-immune response to Influenza A virus (IAV) infection, focusing on the underappreciated role of fibroblasts and their dynamic states. We used 3D quantitative microscopy and scRNAseq to identify IAV-driven fibroblast states, including inflammatory fibroblasts, profibrotic fibroblasts, and adventitial fibroblasts (AFs). Unexpectedly, loss of fibroblast TGFβ signaling led to early enhancement of immuno-modulatory AFs, driving activation of tissue-resident type 2 regulatory T cells (T2-Tregs) and ultimately improving lung functional outcomes. In vitro co-culture systems additionally revealed that AFs act as a niche to support T2-Tregs. Our data suggest an intimate fibroblast-immune crosstalk is required to temporally and spatially balance lung tissue inflammation and repair.

## Introduction

Lung infections caused by common respiratory viruses are major drivers of human morbidity and mortality worldwide. Viral pneumonia poses a severe health risk, with 3-5 million severe cases reported annually worldwide from Influenza infections alone^1^. In both Influenza and COVID-19, acute respiratory distress syndrome (ARDS) is a prevalent complication. ARDS is driven by excessive inflammation caused by diverse lung insults, including pneumonia, trauma, environmental exposures, and systemic sepsis^2,3^. ARDS is characterized by acute lung endothelial and alveolar damage, pulmonary edema, hypoxemia, and multi-organ failure^4^, and has a mortality rate of 25-50%^5,6^. In addition to acute sequelae, lung viral infections are linked to longer-term pulmonary complications, including pulmonary fibrosis, a process correlated with acute disease severity^7,8^. In COVID-19 patients, longitudinal studies identified long-term alterations in diffusion capacity in 40% of patients, with evidence of pulmonary fibrosis^9,10^.

Organ fibrosis is driven by fibroblasts, a type of mesenchymal stromal cell that regulates and supports tissue structure and borders. However, fibroblasts are also increasingly implicated in immune responses, including those occurring within the lung^11,12^. Recent efforts have identified both cross-organ fibroblast states and organ-specific subsets. One conserved profibrotic, myofibroblast state, herein called fibrotic fibroblasts, expresses the highest levels of collagens, α-smooth muscle actin (αSMA), and Cthrc1^+^ (Collagen triple helix repeat containing 1), and, while rare in healthy adults, are a major driver of pulmonary fibrosis^13–17^. In contrast to fibrotic fibroblasts, inflammatory fibroblasts often precede fibrotic fibroblast formation, are associated with acute tissue inflammation, and express Serum amyloid A 3 (Saa3)^18^. Both profibrotic and inflammatory fibroblast states are induced by lung perturbations and reflect discrete, conserved states. Under homeostatic conditions, two major fibroblast states exist in the lung: adventitial and alveolar fibroblasts. Adventitial fibroblasts (AFs) express markers such as Pi16 (Peptidase inhibitor 16), Col14a1 (Collagen type XIV alpha 1), and Ly6a (lymphocyte antigen 6 family member A) and play crucial roles in supporting lung airways, larger vessels, and immune cell subsets^21,22^. In contrast, alveolar fibroblasts specialize in supporting alveolar structure and gas exchange^23,24^.

Just as fibroblast states are heterogeneous, so too are their respective immunological properties. While fibroblasts can express a multitude of cytokines and chemokines upon stimulation, not every fibroblast state possesses the same immune-modulatory profile^25–27^. AFs express the alarmins IL-33 and TSLP (thymic stromal lymphopoietin) and support lymphocytes associated with allergic type 2 inflammation, including ILC2s and Th2s^21^. In contrast, fibrotic fibroblasts are often immunosuppressive, directly suppressing CD8^+^ T cell responses and limiting responses to checkpoint blockade in cancer^28,29^. Within secondary lymphoid organs such as lymph nodes and the spleen, lymphocyte-interactive fibroblasts called fibroblastic reticular cells (FRC) organize T-cell, B-cell, and dendritic cell interactions. Similar structures, named tertiary lymphoid tissue (TLT), also emerge in organs after inflammation. There is a clear spectrum of fibroblast states, each reflecting unique stromal cell positioning, function, and interplay with the immune system.

Regulatory T cells (Tregs) are crucial immunoregulatory lymphocytes in the immune response to influenza due to their suppressive and reparative function^30^. Tregs are major producers of IL-10 and TGFβ, which play prominent roles in local suppression of immune cells^31,32^. Similarly, production of amphiregulin (AREG) by Tregs is critical for the repair of type II alveolar lung epithelial cells (AT2s) during influenza infection. AREG drives repair in other tissues and is specifically produced by tissue-resident type 2-like (T2)-Tregs that express the IL-33 receptor IL1RL1 (ST2), the transcription factor GATA3, and other elements of the type 2 lymphocyte program^35–38^. An IL-33^+^ fibroblast population supports T2-Tregs within visceral adipose tissue (VAT), skeletal muscle, and the oral mucosa. We previously identified an IL-33^+^ AF state in the lung that serves as an immune hub for ILC2 and Th2 cells, suggesting that AFs may also support T2-Tregs^21,39–42^. Here, we show that lung AFs preferentially support T2-Tregs survival and function, serve as critical niche cells in vivo, and that modulation of this niche via TGFβ-signaling ultimately impacts Influenza A viral pneumonia.

## Results

### Lung immune topography in response to influenza infection

First, we determined how lung immune cell positioning (i.e., topography) evolved during Influenza A virus (IAV, strain PR8) infection (**Fig 1A**) using thick-section 3D confocal microscopy. The lung is organized into two distinct regions: distal alveolar (parenchymal) spaces and proximal bronchovasculature. Distal parenchyma is comprised of epithelial-and microvascular-rich alveoli that specialize in gas exchange, whereas proximal bronchovasculature includes large airways (bronchi, bronchioles, αSMA^+^EpCAM^+^), blood vessels (αSMA^+^EpCAM^-^), neuronal innervation, lymphatic drainage (**Fig S1A-B**), and the stromal cells (fibroblasts and mural smooth muscle cells) that support these structures^43,44^. At rest (0 days post-infection/dpi), healthy alveolar type II epithelial cells (AT2s, pro-SPC^+^) were found throughout the distal lung parenchyma, with dispersed, low-level immune cells (CD45^+^) modestly enhanced in the adventitia **(Fig 1B)**. By 7 dpi, immune cells expanded into alveolar areas, accompanied by a concomitant loss of AT2s, consistent with lung damage and loss of functional regions of gas exchange **(Fig 1C)**. By 14 dpi, immune cells reached the distal areas of the lung and encompassed large “damaged patches” **(Fig 1D)**. By 26 dpi, immune cells contracted into altered adventitial spaces with increased TLT structures and with a concomitant increase in alveolar AT2s, indicating lung functional recovery **(Fig 1E)**. Dysplastic epithelial patches, defined by aberrant upper airway epithelial cell hyperplasia (Krt5^+^), were sporadically noted at later times post injury (**Fig S1C**)^45–47^. Lung IAV viral levels, as measured by nucleoprotein transcript (*Np1*), were detectable as early as 3 dpi, peaked at 7 dpi, and were no longer detectable by 9 dpi **(Fig 1F)**. In contrast, lung damage patches continued to enlarge from 7-14 dpi, but subsequently contracted by 26 dpi **(Fig 1G)**.

**Figure 1:**
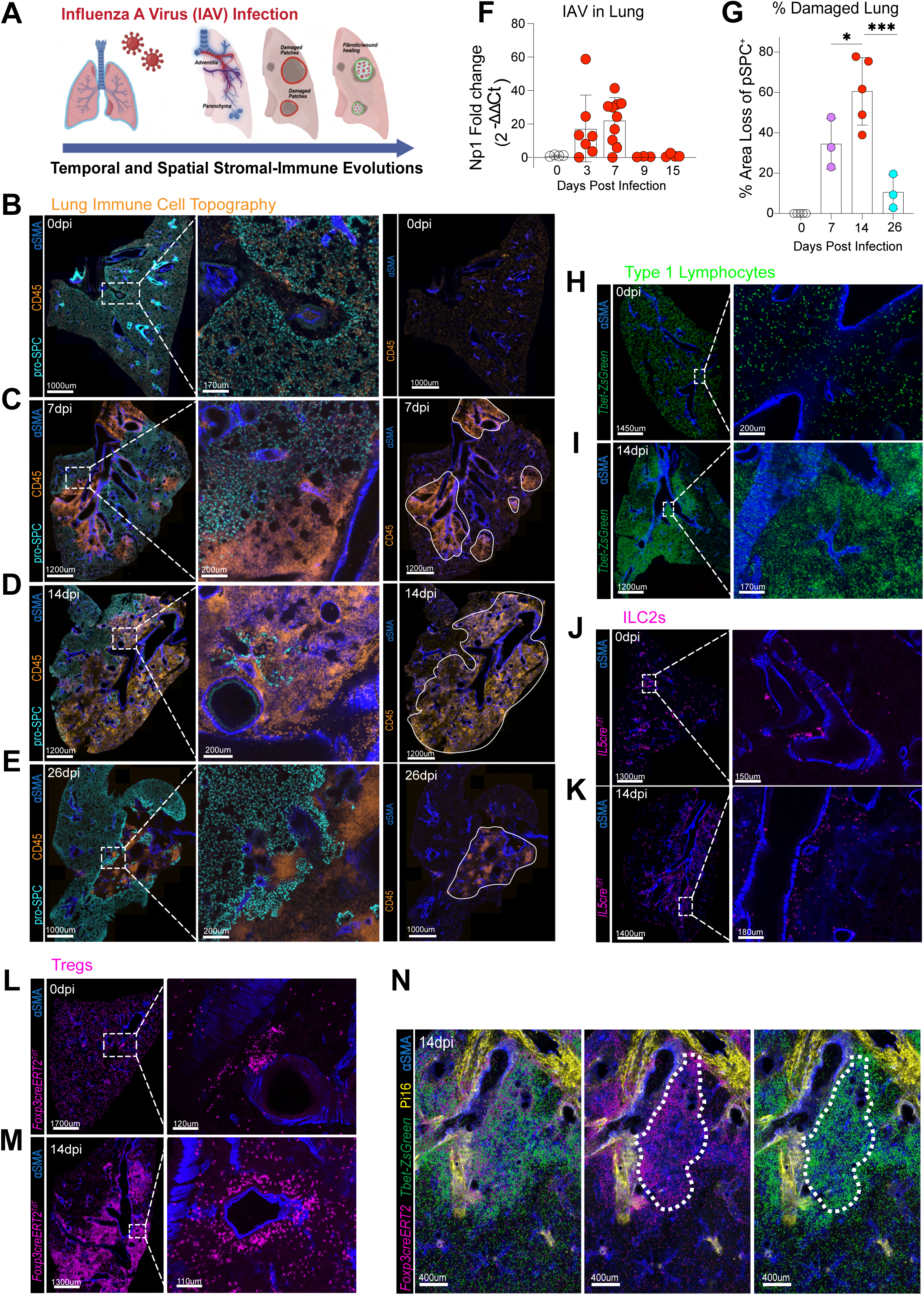
**A,** Schematic illustrating the influenza model of infection in the lung. **B-E**, Time course whole slice and zoomed images of (**B**) 0 dpi, (**C**) 7 dpi, (**D**) 14 dpi, and (**E**) 26 dpi mouse lung stained for AT2s (pro-SPC) and immune infiltrate (CD45). **F**, Time course of viral load using qPCR quantitation of viral nucleoprotein (NP-1). **G**, Time course of relative damage patches (CD45^+^proSPC^-^ regions) in lung slices. **H-I**, Whole slice and zoomed image of T1L (Tbet^ZsGreen+^) localization at (**H**) 0 and (**I**) 14 dpi. **J-K**, Whole slice and zoomed images of ILC2 (IL5^cre^; Rosa26^tdt+^) localization at (**J**) 0 dpi and (**K**) 14 dpi. **L-M**, Thick section and zoomed images of Treg (Foxp3^CreERT2^; Rosa26^RFP+^) localization at (**L**) 0 dpi and (**M**) 14 dpi. **N**, Representative image of Treg (Foxp3^CreERT2^; Rosa26^RFP+^) and T1L (Tbet^ZsGreen+^) localization within a damage patch. **B-E**, **H-M**, 20-40 µM slices. **N,** 200-300µM slices; images represent two or more mice. **G**, Graph shows mean±s.d. *p<0.05, ***p<0.001; one-way ANOVA, Tukey post-test.

Next, we examined how lung lymphocyte subsets and their positioning changed during IAV infection. After early innate- and myeloid cell-focused immune recruitment, influenza drives a dominant anti-viral type 1 immune response, defined by an influx of lymphocytes that produce IFNγ. In uninfected mice, we observed a homogeneous distribution of lung type 1 lymphocytes (Tbet^ZsGreen+^, T1Ls) **(Fig 1H)**, with a majority of T1Ls being intravascular (**Fig S1D**). At 14 dpi, lung damage patches had accumulated T1Ls (**Fig 1I**), many of which were tissue-resident cytotoxic T lymphocytes (ivCD45^-^ CD8^+^ Tbet^+^) **(Fig S1D-F,** 7-14 dpi). Flow cytometry confirmed that tissue-resident IAV-driven lung T1Ls were comprised of NK cells that peaked at 7 dpi and a mix of CD8 T cells and less abundant Th1 CD4^+^ T cells by 14 dpi (**Fig S1G**).

We then sought to determine the positioning of lung type 2 lymphocytes (T2Ls) after IAV infection. T2Ls are traditionally associated with allergic inflammation, are enriched in adventitial fibroblast-dense regions of the proximal lung^48,49^, and have a context-dependent impact on IAV infection outcomes^50–52^. Using a well-validated mouse reporter line, we found effector T2Ls localized to expected lung bronchovascular adventitial regions (IL5^cre^; R26^tdT^ mouse, predominantly identifies ILC2s in resting mice; **Fig 1J)**^21^. By 14 dpi, there was a modest expansion and spread of ILC2s into damaged regions, although they remained largely adventitial-localized **(Fig 1K)**. Treg subset(s) localize closely with ILC2s in the lung^21,53^, and indeed, Tregs were enriched within adventitial spaces (Foxp3^CreERT2^; R26^RFP^ mice; **Fig 1L)**. After IAV infection, Tregs localized either to the lung proximal bronchovasculature (adventitia) or along the borders of alveolar damage patches surrounding dense patches of type 1 lymphocytes **(Fig 1M-N)**. Tregs expanded over time, similar to other helper T cells, peaking at 14 dpi, whereas ILC2s remained relatively stable **(Fig S1H)**. These data suggest that IAV drives distinct spatial and temporal accumulation of type 1 versus type 2 and regulatory lymphocytes.

### Dynamic fibroblast topography in response to influenza infection

Next, we examined lung stromal cell heterogeneity and evolution during IAV infection^54,55^. We found dynamic changes in lung αSMA levels and distribution up to 45 dpi, consistent with altered stromal cells during IAV infection **(Fig S1I-K**). To better parse stromal cell evolution during flu, we first focused on resting lung fibroblast states, confirming that Gli1^+^ fibroblasts (enriched for AFs) were present at the tissue borders, including proximal bronchovasculature and pleural surfaces (Gli1^CreERT2^; R26^tdT^; **Fig 2A**, **Fig 2SA)**. We cross-validated these findings using reporter mice, which accurately identify the spatial positioning of both AFs (Pdgfrα^GFP+^Il33^mcherry+^) and alveolar fibroblasts (Pdgfrα^+^Il33^-^) **(Fig. 2B)**. To further define the topography of lung fibroblast states, we used Xenium single-cell spatial transcriptomics (10X Genomics), validating the predicted localization of AFs and alveolar fibroblasts (**Fig 2C**). Using published single-cell RNA sequencing (scRNAseq) of lung stromal cells **(Fig S2B)**, AFs (Col14a1^+^Pi16^+^) were further parsed into two major subsets, marked by high *Il33* versus high *Aqp1*/*Adh7* **(Fig S2C)**^18^. Using spatial transcriptomics, we found that Aqp1^+^ fibroblasts were preferentially located immediately below the conducting airway epithelial cells and intercalated between airway smooth muscle cells (note that Aqp1 also labels endothelial cells); in contrast, IL-33^+^ AFs (Gli1^+^IL-33^+^) were localized farther from smooth muscle and were abundant around large vessels and, to a lesser extent, outermost regions of airways **(Fig S2D-F),** identifying them as the main immune-modulatory adventitial fibroblasts. We validated these findings via thick-section microscopy **(Fig 2D, Fig S2G-J)**. These data suggest Aqp1^+^ AFs are a subset of cells previously identified as sub-epithelial airway fibroblasts^17,56^, whereas IL-33^+^ AFs are preferentially associated with proximal vasculature and immune cell networks, as we previously described^43^ (**Fig S2K**).

**Figure 2:**
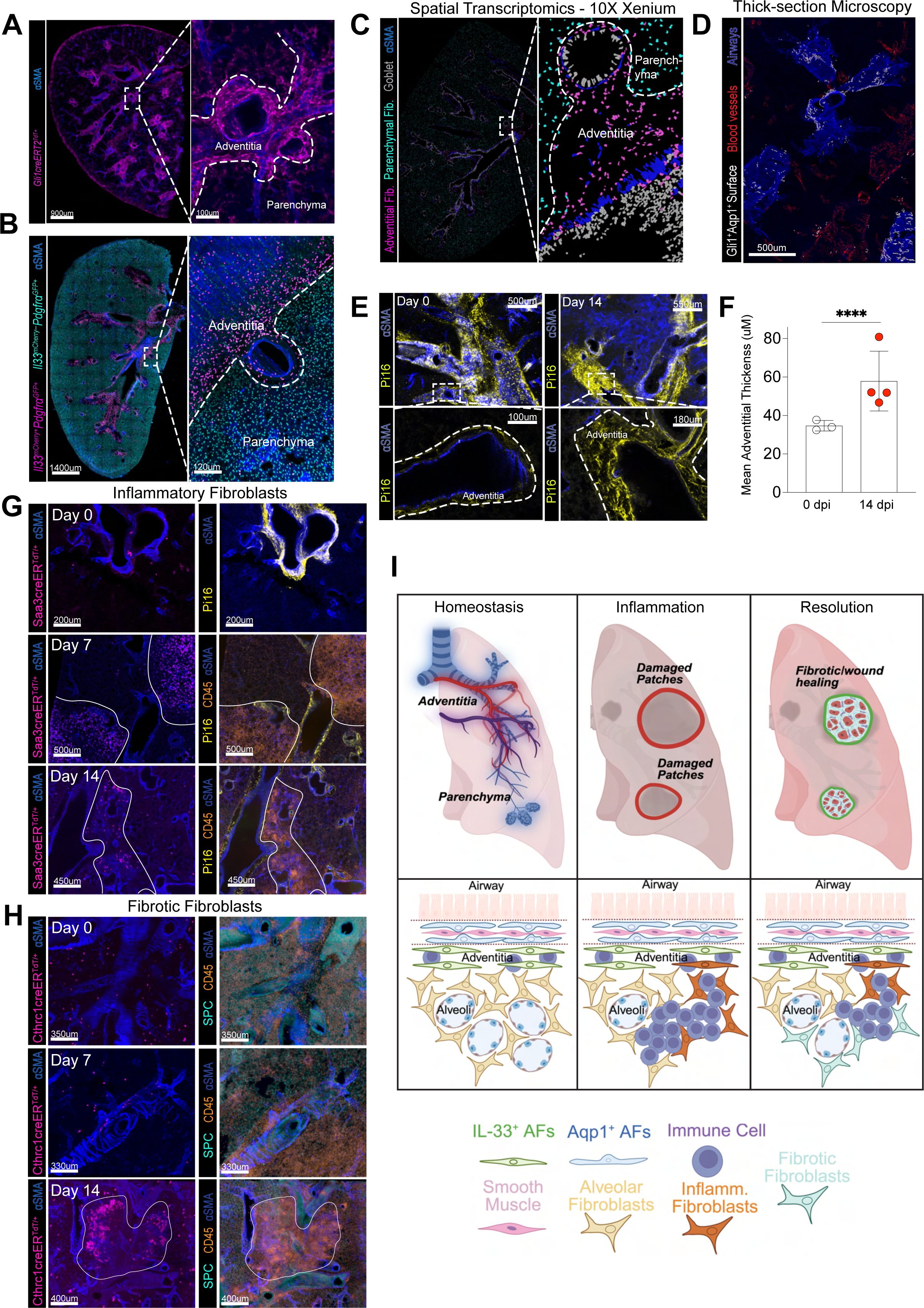
**A**, Representative whole slice and zoomed image detailing adventitial regions (Gli1^+^) labeled by the *Gli1* lineage tracer (Gli1^CreERT2^; Rosa26^tdt^). **B**, Representative whole slice and zoomed image detailing localization of AFs (Pdgfra^GFP+^Il-33^mcherry+^) and alveolar fibroblasts (AlvFs, Pdgfra^GFP+^Il-33^mcherry-^). **C**, Representative whole slice and zoomed image of Xenium spatial sequencing detailing relative localizations of AFs (magenta), AlvFs (cyan), goblet cells (grey), and smooth muscle (blue) cells. **D**, Representative image of Imaris surfaces detailing blood vessels (red) and airway (blue) vasculature in relation to sub-epithelial airway AF surfaces (white). **E**, Thick section and zoomed image of adventitial spaces (white dashed line) marked by Pi16 staining at 0 dpi and 14 dpi. **F**, Imaris quantitation of adventitial thickness from thin section stacks. **G**, Time course images looking at the presence and localization of inflammatory fibroblasts (Saa3^creER^; Rosa26^Ai14+^) at 0, 7, and 14 dpi. Thick-section native imaging of inflammatory fibroblasts (left panels), and adventitial regions (Pi16^+^) and immune cells (CD45^+^, right panels). **H**, Time course images looking at the presence and localization of fibrotic fibroblasts (Cthrc1^creER^; Rosa26^Ai14+^) at 0, 7, and 14 dpi. Thick-section native imaging of fibrotic fibroblasts (left panels), and alveolar regions (pSPC^+^) and immune cells (CD45^+^, right panels). **I**, Schematic illustrating changes in immune-fibroblast population dynamics and localizations over the course of influenza infection. (**A-C,** and **G)**, 20-40µM slices. (**D-E,** and **H)**, 200-300µM slices; images represent two or more mice. (**F**) Graph shows mean±s.d. ****p<0.0001; paired Student’s T-test.

After IAV infection, total adventitial volume expanded by 14 days post-IAV infection (**Fig 2E-F**), resembling AF expansion after lung helminth infection^21^. However, the relative localizations of the IL33^+^ adventitial fibroblasts and Aqp1^+^ sub-epithelial fibroblasts did not change **(Fig S2L-M).** After tissue injury or infection, several conserved and transient fibroblast states emerge, including inflammatory and pro-fibrotic states. Inflammatory fibroblasts (Saa3^CreERT^; R26^tdT^) were largely absent in the lungs of healthy mice. By 7 dpi, they were abundant within alveolar-damaged patches and sparsely present in adventitial regions. By 14 dpi, inflammatory fibroblasts decreased but remained localized to damage patches and the borders of altered adventitia **(Fig 2G)**. To characterize fibrotic fibroblasts, we used the lineage-tracing *Cthrc1* mouse line (Cthrc1^CreERT^; R26^tdT^). Fibrotic fibroblasts were sparse in the resting lung and were minimally induced by 7 dpi. By 14 dpi, fibrotic fibroblasts increased and were found in alveolar damaged patches, often at the interface of alveolar or adventitial regions (**Fig 2H**). Together, these data demonstrate lung fibroblast temporal and spatial heterogeneity after IAV infection, with the emergence of earlier inflammatory and later fibrotic fibroblast states within damaged lung patches, a pattern resembling changes observed in models of lung fibrosis^18^ (**Fig 2I**).

### Loss of fibroblast TGFβ signaling improves lung health during influenza infection

Lung fibrosis is a complication of severe respiratory virus infection^8,57,58^, suggesting fibrotic fibroblasts may functionally contribute to lung remodeling after IAV infection. TGFβ is a key driver of the fibrotic fibroblast state in the lung, brain, and other tissues^59,20^. While an excess of fibrotic fibroblasts can drive fibrosis, a complete loss of this state can also be detrimental to organ functional recovery^17,20^. To test the functional role of fibrotic fibroblasts in IAV infection, we inducibly deleted TGFβ signaling within all fibroblasts (**Fig 3A**), hereafter called TGFbr2 conditional knockout mice (TGFbr2 cKO, Col1a2^CreERT2^; Tgfbr2^F/F^ mice) and infected mice with IAV. It was previously observed that inducible deletion of fibroblast TGFβ signaling in adult TGFbr2 cKO mice did not impact animal health at rest^20^. However, after IAV infection, TGFbr2 cKO mice showed unexpectedly improved lung and overall health as early as 7 dpi, reflected by increased blood oxygen levels and body weight (**Fig 3B,C**). Viral titers at 3 dpi were not altered in TGFbr2 cKO mice, suggesting altered lung health was not directly linked to alterations in viral clearance **(Fig S3A)**.

**Figure 3:**
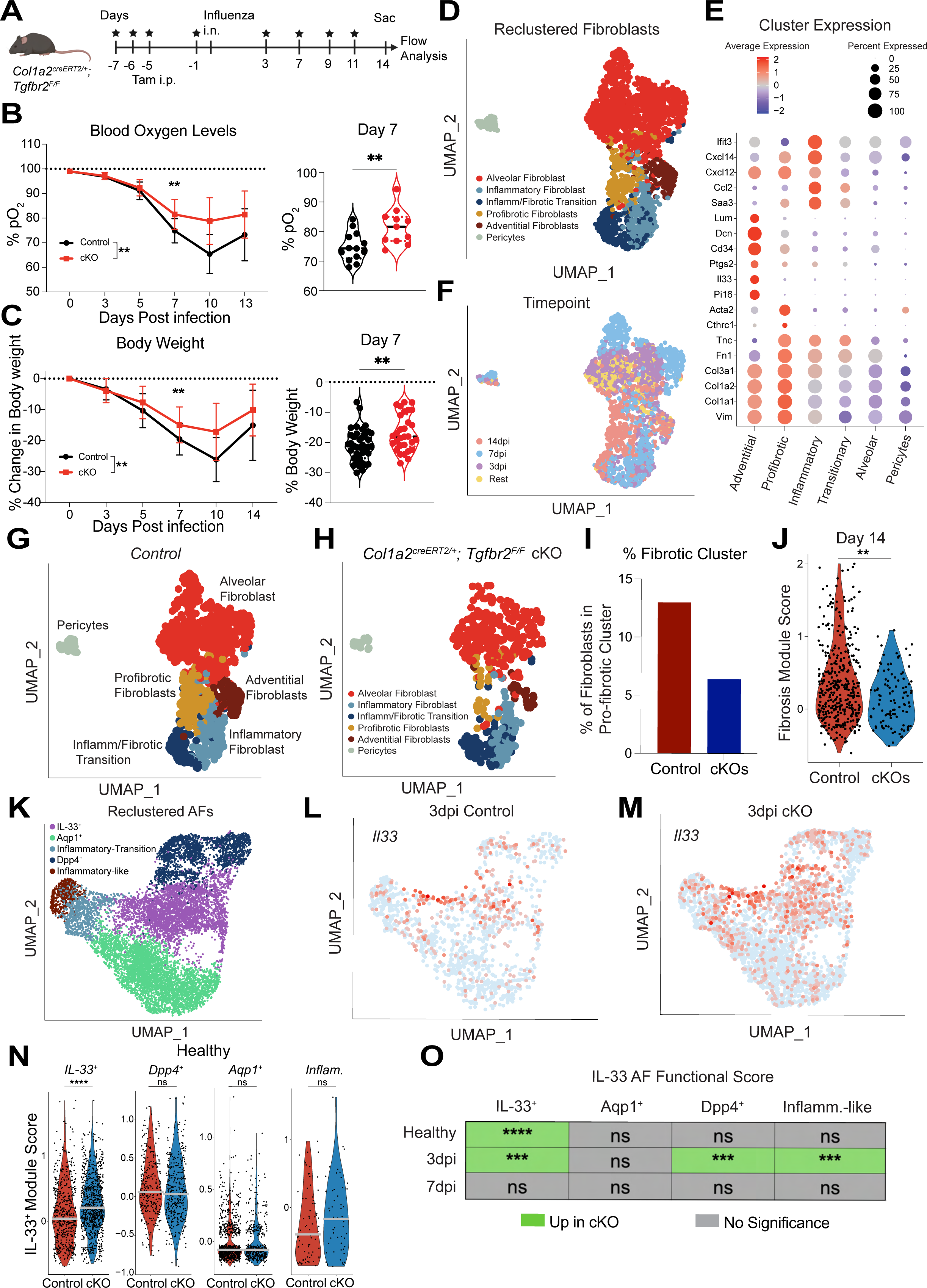
**A**, Graphical diagram illustrating mouse model and timing of tamoxifen injections (i.p.), influenza inoculation (i.n.), and sacrifice. **B**, Pulse oximetry time course in controls/TGFbr2 cKOs (left). Pulse oximetry measurements at 7 dpi comparing controls and TGFbr2 cKOS (right).**C**, Body weight time course in controls/ TGFbr2 cKO mice (left). Body weight measurements at 7 dpi comparing controls and TGFbr2 cKO mice (right). **D**, UMAP of fibroblast clusters showing various fibroblast states, including alveolar, inflammatory, “inflammatory-fibrotic” transition, profibrotic, adventitial, and pericytes. **E**, Dot plot showing marker genes for all cellular clusters. **F**, UMAP of fibroblast clusters illustrating timepoint metadata. **G-H**, UMAP of respective genotype fibroblast clusters comparing (**G**) controls to (**H**) TGFbr2 cKOs. **I**, Quantitation of normalized proportion of the fibrotic fibroblast cluster between control (red) and TGFbr2 cKO (blue) mice. **J**, Fibrosis module score (upregulated genes that are associated with a pro-fibrotic phenotype) on total fibroblast cluster between control and TGFbr2 cKO mice at 14 dpi. **K**, UMAP of reclustered adventitial fibroblasts illustrating various AF states, including immune-modulatory IL-33^+^, sub-epithelial airway Aqp1^+^, fascial-like Dpp4^+^, inflammatory-transition, and inflammatory-like. **L-M**, UMAP plots showing *Il33* gene expression at 3 dpi in (**L**) control and (**M**) TGFbr2 cKO AFs. **N**, IL-33 functionality scores (genes associated with immune-modulatory capacity of IL-33^+^ AFs) applied across the following AF states: immune-modulatory, fascial-like Dpp4^+^, airway-associated, and inflammatory-like at 3 dpi. **O**, Table illustrating IL-33 functionality comparisons across 0, 3, and 7 dpi. Graphs show mean±s.d. ns = not significant, **p<0.01, ***p<0.001, ****p<0.0001; Mixed-effect analysis with Geisser-Greenhouse correction (**B-C, left panels**); Student’s T-test (**B-C, right panels**); Mann-Whitney test (**J, N-O**).

To define the cellular dynamics underpinning lung functional improvements in TGFbr2 cKO mice, we performed lung single-cell RNA sequencing (scRNAseq) across the IAV infection time course (0, 3, 7, and 14 dpi), sorting and enriching for fibroblasts and T cells (**Fig S3B)**. Unsupervised clustering identified 6 broad cell subsets: lymphocytes, myeloid cells, stromal cells, endothelial cells, epithelial cells, and doublets **(Fig S3C,D)**. Sub-clustering on stromal cells identified several subsets, including alveolar fibroblasts (AlvF), inflammatory fibroblasts, inflammatory/fibrotic transition fibroblasts, fibrotic fibroblasts, adventitial fibroblasts (AF), and mural cells (pericytes/smooth muscle cells) (**Fig 3D**), resembling findings after lung bleomycin-induced fibrotic injury^17,18^. AFs expressed expected markers including *Pi16, Il33, Dcn, and Cd34*; inflammatory fibroblasts expressed *Saa3, Ccl2, Ifit3, and Cxcl14;* fibrotic fibroblasts expressed profibrotic markers, including *Cthrc1, Acta2, Fn1, and Tnc* (**Fig 3E**). Consistent with confocal microscopy, inflammatory fibroblasts were present by 3 dpi, most prominent at 7 dpi, and decreased by 14 dpi, whereas fibrotic fibroblasts were most prominent at 14 dpi; in contrast, both subsets were largely absent at rest (**Fig 3F**).

Innate immune cells, such as neutrophils and NK cells, are critical early responders in influenza infection^60–62^. Therefore, we investigated potential changes in both myeloid cells and innate lymphoid cells after IAV in our TGFbr2 cKO animal scRNAseq. Within myeloid populations, we identified dendritic cells (DCs), neutrophils, Arg1^+^ macrophages, monocytes, and other myeloid cells **(Fig S3E-G)**. Results showed small differences in the relative proportions of myeloid cell subsets at both 3 dpi and 7 dpi **(Fig S3H-I)**. Flow cytometry did not identify consistent alterations in PMN or NK cell number (see below, **Fig S4A-D**). These data, along with comparable viral titers among TGFbr2 cKO and control mice at 3 dpi, suggest that the observed lung health differences were not due to altered viral clearance or neutrophil- or NK cell-mediated inflammation.

Next, we examined how TGFβ-signaling affected IAV-driven lung fibroblasts. As expected, by 14 dpi TGFbr2 cKO mice had reduced fibrotic fibroblasts and a fibrosis module score (**Fig 3G-J**). Although the diminution of fibrotic fibroblasts in TGFbr2 cKO mice could, in theory, drive improved lung health, fibrotic fibroblasts did not develop until later time points (∼14 dpi), which did not correlate with improved animal health as early as 7 dpi. This suggested that TGFbr2 cKO mice may have other changes in fibroblast subsets, including potential shifts in AF states at earlier time points post-IAV infection.

To better parse AF subsets after IAV infection, we performed a second scRNAseq experiment, enriching for stromal cells from the lungs of control and TGFbr2 cKO mice at early time points post-IAV (0 dpi, 3 dpi, and 7 dpi). Unbiased clustering revealed 7 distinct stromal cell states: Adventitial fibroblasts (AF), alveolar fibroblasts (AlvF), inflammatory fibroblasts, pericytes, vascular- and peribronchial- smooth muscle cells (SMC), and mesothelial cells **(Fig S3J-K)**. Unbiased sub-clustering was conducted on the AF cluster **(Fig 3K)**, identifying expected Aqp1^+^ sub-epithelial airway AFs and IL-33^+^ AFs, as well as Saa3^+^ inflammatory AFs, and a Dpp4^+^ AF state with similarities to fascial fibroblasts in the skin (**Fig S3L-O**)^63^ but distinct from mesothelial cells **(Fig S3P)**. By 3 dpi, AF subset proportions were similarly abundant between cKO and control mice; however, cKO mice displayed higher *Il33* gene expression across multiple AF subsets **(Fig 3L-M)**.

Furthermore, an IL-33^+^ AF-derived module score (genes include *Il33, Ptgs2, Ccl11,* and other AF-associated genes^17,21,64^) was applied to the various AF subsets, which showed a higher score in the TGFbr2 cKO mice in the IL-33^+^ AF state at rest **(Fig 3N)**; by 3 dpi, this module was higher in TGFbr2 cKO mice across multiple AF states, although these differences were no longer present by 7 dpi **(Fig 3O)**. These data suggest that loss of fibroblast TGFβ signaling increased IL-33^+^ AF immune-modulatory functionality at rest and early time points after IAV, potentially underpinning the functional improvements in IAV-infected cKO mice. More broadly, these findings suggest that TGFβ signaling in fibroblasts exerts dual roles, driving the fibrotic fibroblast state but also directly restricting the IL-33^+^ AF immunomodulatory state and function.

### Type 2-like Tregs localize to the adventitial niche and expand in cKO mice after IAV infection

IL-33^+^ AFs perform multiple functions, including maintenance and remodeling of the adventitia, with similarities to IL-33^+^ fascial fibroblasts at other natural tissue borders. IL-33^+^ AFs also regulate ILC2 and Th2 allergic lymphocytes that drive type 2 inflammation^21,65,66^. Lung ILC2s co-localize near Tregs and, during type 2 inflammation, support Treg subset(s) that express portions of the “type 2 lymphocyte” program, including the transcription factor Gata3 and the IL-33 receptor ST2 (IL1RL1)^53,67^. As such, we hypothesized that the increase in IL-33^+^ immunomodulatory AFs in TGFbr2 cKO mice may alter lung Tregs or other adventitial immune cells, thereby contributing to improved lung health after IAV infection. Indeed, using flow cytometry, we compared the immune compartment of TGFbr2 cKO mice to littermate age-matched controls during IAV infection, and found tissue-resident type 2 (T2) Tregs were elevated in cKO mice at 3 dpi **(Fig 4A-B)**, with increased proliferation **(Fig 4C-D)**. We observed few differences in frequencies of other immune cell subsets at rest, 3, 7, and 14 dpi, including: tissue resident CD8^+^ T cells, ψ8 T cells, ILC2s, macrophages, dendritic cells (DC1, DC2s), alveolar macrophages, monocytes, neutrophils, eosinophils, and NK cells **(Fig S4A)**, consistent with our scRNAseq data. Moreover, few differences were observed in frequencies of specific subsets of tissue Type 1 Tregs and Type 1/2 CD4 effector cells (**Fig S4B**). Little to no differences were observed between genotypes in the numbers of the above-mentioned immune cells (**Fig SC-D**).

**Figure 4:**
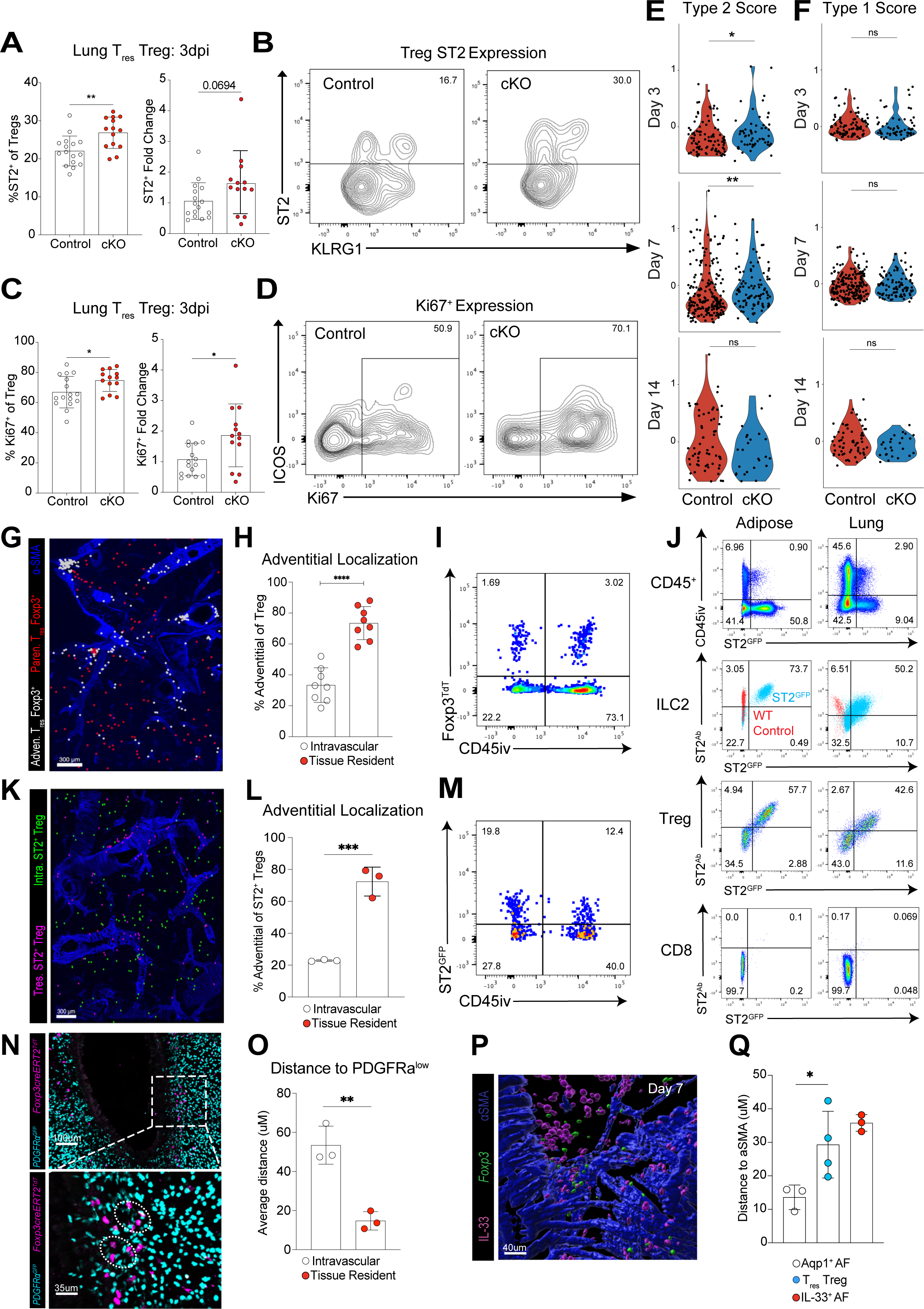
**A**, Quantitation comparing frequency (left panel) and numbers (right panel) of type 2-like (ST2^+^) tissue resident Tregs between control (white) and TGFbr2 cKO (red) mice. **B**, Representative FACS plot showing differences in ST2 and KLRG1 expression in tissue resident Tregs between control and TGFbr2 cKO mice. **C**, Quantitation comparing frequency (left panel) and numbers (right panel) of proliferating (Ki67^+^) type 2-like tissue resident Tregs between control (white) and TGFbr2 cKO (red) mice. **D**, Representative FACS plot showing differences in Ki67 expression in type 2-like tissue resident Tregs between control and TGFbr2 cKO mice. **E-F**, Module scores looking at type 2 (**E**) and type 1 (**F**) characteristics of the Treg population across 3, 7, and 14 dpi. **G**, Thick-section image illustrating Imaris surfaces of adventitial (white) and parenchymal (red) tissue-resident Tregs. **H**, Quantitation comparing proportions of tissue resident and parenchymal Tregs that are adventitial-associated (<20µM from αSMA). **I**, Representative FACS plot validating CD45i.v. staining. **J**, FACS plots illustrating the specificity of ST2 expression of the ST2^GFP^ reporter across various immune cells. **K**, Thick-section image illustrating Imaris surfaces of tissue resident type 2 Tregs (magenta) and intravascular type 2 Tregs (green). **L**, Quantitation comparing proportions of tissue resident and intravascular Type 2-like Tregs that are adventitial-associated (<20µM from αSMA). **M**, Representative FACS plot validating ST2^GFP^ expression within Tregs. **N**, Representative thick section and zoomed image of native imaging looking at relative localizations of AFs (PDGFRα^low^) and Tregs (Foxp3^CreERT2^; Rosa26^RFP+^). **O**, Imaris quantification of average distance of intravascular (IV^-^) and tissue resident (IV^+^) Tregs to PDGFRα^low^ AFs. **P**, Zoomed image of Imaris surfaces looking at relative localizations of Tregs (green) and IL-33^+^ cells (magenta). **Q**, Imaris quantitation of average distance of Aqp1^+^ AFs, IL-33^+^ AFs, and tissue resident Tregs to αSMA. (**G, K, N, P**) 200-300µM slices; images represent two or more mice. Graphs show mean±s.d. ns = not significant, *p<0.05, **p<0.01, ***p<0.001, ****p<0.0001; Student’s T-test (**A, C, H, L, O**); Mann-Whitney test (**E-F**). One-way ANOVA, Tukey post-test (**Q**).

Next, we used our scRNAseq data to cross-validate the potential elevation of lung T2-Tregs in cKO IAV-infected mice. Within the CD4^+^ T cell cluster, we identified a small Treg cluster and isolated it for further analysis **(Fig S4E**, clusters 0, 1, and 5). A T2-Treg module score (genes including *Gata3, Il1rl1, Areg, Setd2, Il2ra*) was elevated at both 3 dpi and 7dpi in TGFbr2 cKO Tregs as compared control Tregs **(Fig 4E)**. A Treg “type 1 immune module” score (including *Tbx21, Cxcr3, Cx3cr1*) showed no differences across all time points post IAV infection, consistent with our flow cytometry data **(Fig 4F)**. We conclude that T2-Tregs are preferentially activated early post-IAV infection in TGFbr2 cKO mice, coinciding with increased functionality of the immunomodulatory IL-33^+^ AF state.

To further validate a Treg-AF relationship, we examined the localization of lung Tregs using thick-section confocal microscopy in Treg-lineage-tracing mice (*Foxp3^CreERT2^; R26^RFP^*). We utilized intravenous labeling (IV) with either CD45 or Thy1.2 antibodies to determine tissue residency. Tissue-resident Tregs (IV^-^) localized preferentially to the lung adventitia, whereas intravascular Tregs were broadly distributed in the alveolar microvasculature **(Fig 4G-H, Fig S4F)**, resembling NK cells (Tbet^+^ CD3ε^-^) that are typically intravascular in unperturbed mice^68^. Using flow cytometry, we validated the presence of a minor subset of tissue-resident Tregs in naïve mouse lungs **(Fig 4I)**. Next, we parsed Treg states to identify type 2 (T2) or type 1 (T1) Tregs. We used a novel mouse reagent that marks cells expressing the IL-33 receptor ST2 (IL1RL1) with GFP and validated the expected GFP expression in ST2^+^ type 2 lymphocytes **(Fig 4J, Fig S4G)**. T2-Tregs (*Foxp3^creERT2^; R26^RFP+^; ST2^GFP+^*) were enriched within the tissue-resident lung fraction and preferentially localized to the lung adventitia (**Fig 4K-M)** and in proximity to putative ILC2s (ST2GFP^+^; FoxP3^-^) **(Fig S4H)**.

The temporal landscape of lung Treg subsets is dynamic after flu infection, and we confirmed with flow cytometry that T2-Tregs were the most abundant early post-infection, whereas T1-Tregs peaked by 14 dpi **(Fig S4I-J),** similar to published results^33,48^. To test Treg localization relative to fibroblasts, we used PDGFRα^GFP^; *Foxp3^creERT2^; R26^RFP^* mice. As expected, tissue-resident Tregs localized tightly with PDGFRα^GFP-lo^ AFs in adventitial regions **(Fig 4N-O, Fig S4K)**. Additionally, Tregs were more closely associated with immunomodulatory AFs (IL-33^+^) compared to airway sub-epithelial AFs (Aqp1^+^) **(Fig 4P-Q)**. Together, these data suggest an early activation of adventitial-localized T2-Tregs in cKO mice, which may explain the enhanced lung health outcomes, particularly given the described beneficial roles of T2-Tregs in promoting epithelial repair through secretion of Areg, production of IL-10, and local suppression of hyperactive immune cells^30,34,69,70^.

### Lung adventitial fibroblasts directly and preferentially support T2-Tregs

Given their spatial and temporal association, we predicted that lung IL-33^+^ AFs may cross-regulate T2-Tregs. To better understand this putative stromal-immune interaction, we developed a co-culture system using primary lung AFs and Tregs. AFs were sort-purified (CD45^-^CD31^-^Pdgfra^+^Sca1^+^) and grown to confluence over 7 days, after which Tregs were sort-purified from the spleen and lymph nodes of Foxp3^GFP^ reporter mice, and added to AFs with or without exogenous TCR stimulation (**Fig 5A**). Tregs cultured with AFs had higher viability, proliferation, and markers of a T2-state (ST2^+^KLRG1^+^), resembling Tregs treated with TCR stimulation, and increased compared with media alone controls (**Fig 5D-G**). In contrast to AFs, lung alveolar fibroblasts (CD45^-^CD31^-^Pdgfra^+^Sca1^-^) induced less Treg proliferation **(Fig S5A)**. When Tregs were grown in conditioned media from AFs, their proliferation remained low, similar to Treg-alone conditions, albeit with a modest increase in T2-Treg markers (ST2^+^KLRG1^+^) (**Fig 5E-F**). AF-conditioned media was sufficient to support Treg viability (**Fig 5G**), similar to previous results demonstrating that AF-derived soluble factors can support ILC2s. Next, we co-cultured Tregs with AFs in transwell plates and found reduced Treg proliferation and viability (**Fig S5B-D**). Together, these data indicate that AFs can promote Treg viability, proliferation, and “type 2” characteristics, acting primarily through contact-dependent or local mechanisms, although AF-derived soluble signals may provide additional supportive signals.

**Figure 5:**
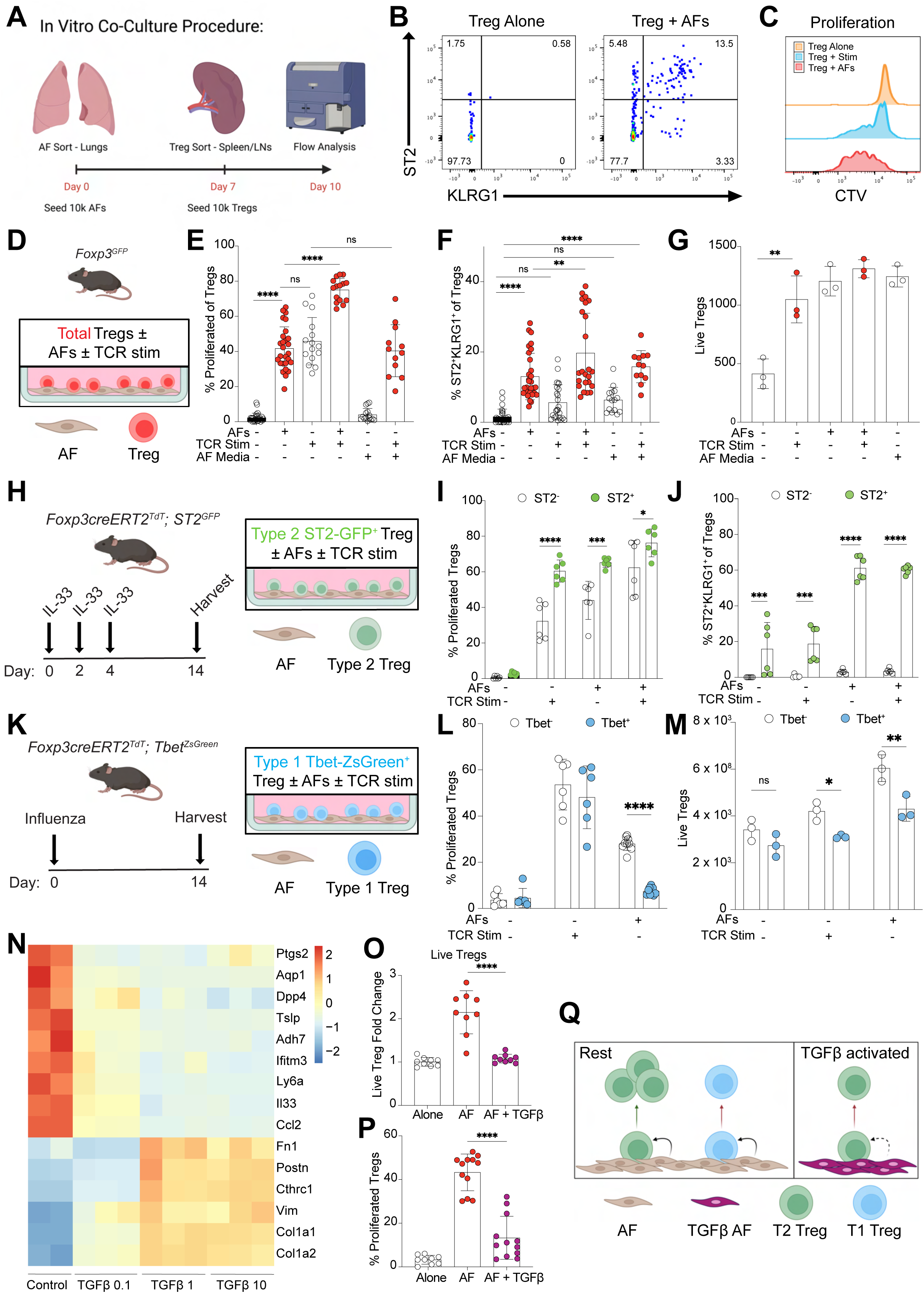
**A**, Graphical illustration of *in vitro* co-culture procedure. **B**, Representative FACS plot showing the effect of AFs on ST2 and KLRG1 expression on Tregs. **C**, Representative plot showing effect of AFs on Treg proliferation (CTV staining). **D**, Graphical illustration illustrating total spleen and nodal Tregs sorted for culture from Foxp3^eGFP^ mice. **E-G**, Quantitation comparing the effects of a combination of AFs, stim (CD3/CD28 beads), and AF conditioned media on Treg (**E**) proliferation, (**F**) ST2^+^KLRG1^+^ expression, and (**G**) Treg viability. **H**, Graphical illustration of IL-33 injection timeline in Foxp3^CreERT2^; ST2^GFP^ mice to sort out type 2 Tregs. **I-J**, Quantitation comparing the effects of a combination of AFs and stim on ST2^+^ and ST2^-^ Treg (**I**) proliferation and (**J**) ST2^+^KLRG1^+^ expression. **K**, Graphical illustration of infection timeline in Foxp3^CreERT2^; Tbet^ZsGreen^ mice to sort out type 1 Tregs. **L-M**, Quantitation comparing the effects of a combination of AFs and stim on Tbet^+^ and Tbet^-^ Treg (**L**) proliferation and (**M**) viability. **N**, Heatmap illustrating markers associated with AFs and fibrotic fibroblasts across a dose titration of TGFβ treatment. **O-P**, Quantitation comparing effects of AFs and TGFβ-treated AFs on Treg (**O**) viability and (**P**) proliferation. **Q**, Graphical illustration of the interactions between AFs and TGFβ-treated AFs on type 2 and type 1 Tregs. Graphs show mean±s.d. ns = not significant, *p<0.05, **p<0.01, ***p<0.001, ****p<0.0001; one-way ANOVA, Tukey post-test (**E-G, O-P**); two-way ANOVA, Sidak’s post-test (**I-J**, and **L-M**).

A traditional source of support is through Treg T cell receptor (TCR) signaling via MHC class II from antigen-presenting cells, which could include AFs. However, purified AFs from MHCII^-/-^ mice were similarly able to support T2-Tregs in co-culture **(Fig S5E**). AFs are also physiologic producers of IL-33, which we and others have shown supports type 2 lymphocytes^21,66^. To test how IL-33 may impact Treg support, we first added exogenous IL-33 to the co-culture system. Although IL-33 by itself could not support Tregs, IL-33 synergized with AFs to better support T2-Tregs **(Fig S5F-I)**. To further test the role of IL-33, wild-type and IL-33^-/-^ AFs were sorted and found to support Tregs similarly (**Fig S5J-K**), consistent with our previous work showing that AFs do not release IL-33 in vitro. Together, these data indicate that while IL-33 is not required, it can work in concert with other signals from AFs to amplify T2-Tregs.

Next, we sorted-purified effector Tregs (CD44^+^CD62L^-^) and naive Tregs (CD44^-^CD62L^+^) and cultured them with AFs, finding that effector Tregs showed higher proliferation and T2-like (ST2^+^KLRG1^+^) marker expression than their naive counterparts **(Fig S5L-M)**. To further test AF support for Treg subsets, we sorted ST2^+^ T2-Tregs and ST2^-^ Tregs from IL-33-injected mice for co-culture **(Fig 5H)**. We found that ex vivo T2-Tregs (ST2^+^) had higher proliferation and viability than their ST2^-^ counterparts (**Fig 5I-J, Fig S5N**). However, AFs did not induce T2-Treg gene expression (ST2^+^KLRG^+^) from purified ST2^-^Tregs. To further delineate if there was preferential AF support for T2-Tregs as compared to T1-Tregs, we purified Tbet^+^ and Tbet^-^ Tregs from the lungs of IAV-infected mice at 14 dpi and used them in co-culture with AFs **(Fig 5K)**. T1-Tregs (Tbet^+^) had reduced proliferation and viability when cultured with AFs, compared to Tbet^-^ Tregs (**Fig 5L-M**). We conclude that AFs preferentially support pre-existing, or poised, effector T2-Tregs.

Universal fibroblasts were originally identified as a cross-tissue subset with increased mesenchymal cell plasticity^22^. AFs and similar border/fascial fibroblasts are enriched for this universal fibroblast state, and can adopt fibrotic fibroblasts, inflammatory fibroblast, and lymphocyte-interactive (FRC-like) fibroblast states. We first investigated whether AFs could adopt an inflammatory fibroblast state in vitro. We conducted RNA sequencing on sorted AFs cultured with IL-1β, a signal known to drive this state^18^. We found that AFs cultured with IL-1β had increased mRNA levels for genes associated with inflammatory fibroblasts, including *Saa3* and *Ccl2* **(Fig S5O-P).** Next, we investigated whether AFs could adopt a fibrotic fibroblast state. We cultured AFs at varying TGFβ concentrations and performed bulk RNA sequencing. AFs showed a dose-dependent response to TGFβ, with higher doses leading to increased expression of fibrosis-related genes, including *Cthrc1, Fn1, Col1a1,* and *Col1a2;* however, all TGFβ doses resulted in a loss of AF-associated immunomodulatory signature genes, including *Il33, Ptgs2, Tslp,* and *Ccl11* **(Fig 5N, Fig S5Q-R)**. To test whether TGFβ signaling directly impacted the support of AFs for Tregs in vitro, we pre-treated AFs with TGFβ before seeding with effector Tregs (CD44^+^); AFs’ ability to support Treg proliferation and viability was reduced (**Fig 5O-P**), with a trend in reduction in T2-Treg markers (ST2^+^KLRG1^+^) **(Fig S5S)**. Overall, these data suggest that AFs exposed to a range of TGFβ concentrations downregulate the AF immunomodulatory program, whereas higher concentrations of TGFβ are required to drive a profibrotic program **(Fig 5Q)**.

### Loss of T2-Tregs leads to exacerbated lung health in IAV infection

To determine the functional role T2-Tregs may play during influenza infection, we used T2-Treg-deleter mice (Foxp3^CreERT2^; R26^RFP^; Gata3^F/F^). Of note, non-inducible versions of this mouse strain have been previously used in other mouse models^34,71^, although Treg Gata3 expression during thymic development may have influenced these phenotypes. We first validated the mouse model by injecting tamoxifen intraperitoneally and analyzing lungs 14 days later **(Fig. 6A)**. As expected, we saw a specific and sensitive loss of T2-Tregs marked by Gata3, ST2, and KLRG1 **(Fig 6B-F)**, without overt signs of mouse autoimmunity. In contrast, no differences in frequencies or total numbers of tissue resident Tregs, CD4^+^ effector, and CD8^+^ effector T cells were observed (**Fig S6A-C**), suggesting a specific impact on T2-Tregs. After IAV infection, T2-Treg deleter mice showed consistent loss of T2-Tregs from 3 dpi to 14 dpi (**Fig 6G-H**). Loss of T2-Tregs led to exacerbated lung health, as measured by both peripheral blood oxygen levels and body weight **(Fig 6I-J)**. Other immune cell populations, including tissue-resident CD8^+^ T cells, tissue-resident type 1-like Tregs, αβ T cells, ILC2s, macrophages, alveolar macrophages, monocytes, neutrophils, eosinophils, and NK cells, were not different between the groups (**Fig. S6D-N**). These data show that T2-Tregs play a significant role in protecting lung health during IAV infection and suggests that elevated T2-Tregs in TGFβr2 cKO mice are responsible for improved lung function (**Fig. S6P**).

**Figure 6:**
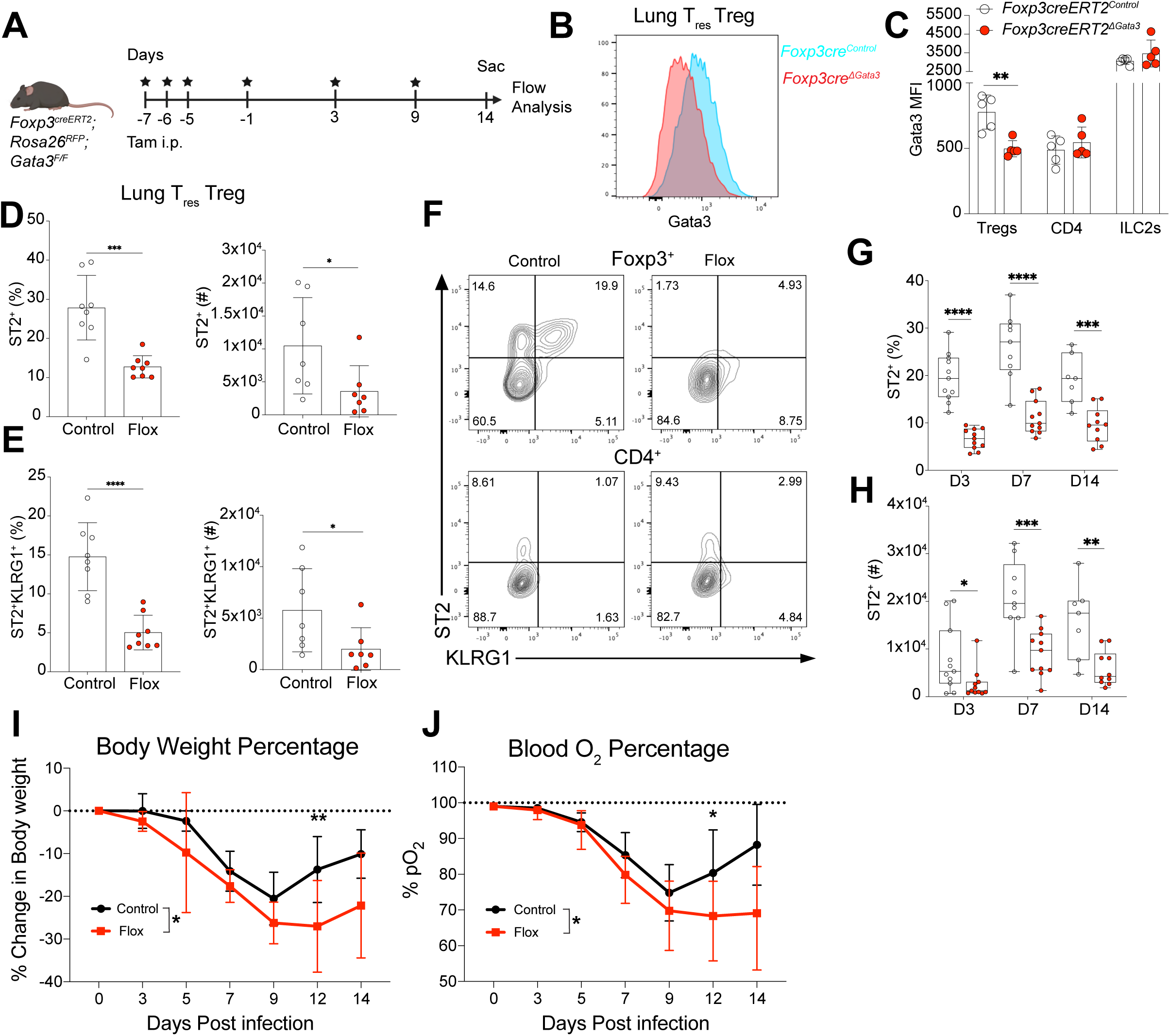
**A**, Graphical illustration showing timeline of tamoxifen injections in T2 Treg deleter mice. **B**, Graph showing differences in Gata3 MFI between control and T2 Treg deleter mice. **C**, Quantitation showing differences in Gata3 MFI across Tregs, CD4 effector cells, and ILC2s. **D**, Quantitation measuring frequency and numbers of ST2^+^ Tregs between control and flox mice. **E**, Quantitation measuring frequency and numbers of ST2^+^KLRG1^+^ Tregs between control and flox mice. **F**, Representative FACS plots illustrating ST2 and KLRG1 expression in Tregs and CD4^+^ T cells in control and T2 Treg deleter mice. **G-H**, Quantitation at 3, 7, and 14 dpi showing continuous loss of ST2 expression in both (**G**) frequency and (**H**) numbers in Tregs. **I-J**, Time course graphs illustrating exacerbated (**I**) peripheral blood oxygen and (**J**) body weights in flox mice. Graphs show mean±s.d. *p<0.05, **p<0.01, ***p<0.001, ****p<0.0001; one-way ANOVA, Tukey post-test (**C**). Student’s T-test (**D-E, G-H** (for each time point)). Mixed-effect analysis with Geisser-Greenhouse correction (**I-J**).

## Discussion

Here, we uncovered a functionally critical lung adventitial fibroblast – T2-Treg niche. Increased functionality of immunomodulatory AFs, driven by reduced TGFβ signaling, led to early activation and expansion of type 2-like (T2) tissue Tregs, ultimately improving lung health in a mouse model of viral influenza. Fibroblasts are heterogeneous and functionally malleable, with recent work delineating diverse, conserved fibroblast states including (1) *Cthrc1^+^* profibrotic fibroblasts, (2) *Pi16^+^*adventitial/border fibroblasts, (3) *Saa3^+^* inflammatory fibroblasts, and (4) *Ccl19*^+^ lymphocyte-interactive fibroblasts that resemble fibroblastic reticular cells (FRC) in the lymph nodes and spleen. Each fibroblast state plays a unique role in local tissue homeostasis, injury, repair, and disease^17,18,43,72^. While tools such as scRNAseq have helped uncover fibroblast heterogeneity, the state-specific spatial-temporal positioning, regulation, and functional contributions of fibroblasts remain unclear. Using a combination of genetic tools, scRNAseq, and thick-section 3D confocal imaging, we defined the fibroblast state positioning within the lung at rest and after Influenza A virus infection. Saa3^+^ inflammatory fibroblasts developed early after IAV infection and were predominantly confined to damaged lung regions. Profibrotic Cthrc1^+^ fibroblasts developed later during influenza infection, were dependent on TGFβ signaling, and were located at the interface between damaged and healthy lung tissue. We also found unexpected heterogeneity in adventitial fibroblasts after viral infection. These include validation of sub-epithelial airway-associated AFs (Aqp1/Adh7+), perivascular IL-33^+^ immunomodulatory AFs, and the emergence of IAV-driven inflammatory (Saa3^+^) AF states that localize to the adventitial borders. In addition to the role of TGFβ in promoting fibrotic fibroblasts, we uncovered a secondary role in limiting IL-33^+^ AFs. Preventing TGFβ signaling in immunomodulatory AFs preserved their functional capacity during IAV infection, leading to increased T2-Tregs and improved viral pneumonia outcomes.

Our findings highlight the role of immunomodulatory AFs in supporting tissue-resident T2-Tregs. T2-Tregs are a Treg state that is enriched within resting tissues and can play beneficial roles (e.g., wound repair) or pathologic roles (e.g., cancer). Thick-section microscopy revealed that lung T2-Tregs localize preferentially to adventitial spaces near immunomodulatory IL-33^+^ AFs. Co-culture experiments demonstrated direct, preferential support by AFs for T2-Tregs, which synergizes with IL-33 signaling. Although AFs in vitro did not release IL-33, they can be relevant in vivo sources of IL-33, and this pathway likely represents one mechanism by which AFs regulate T2-Tregs. How AFs support T2-Treg survival and proliferation remains to be elucidated, but involves contact-dependent mechanisms, potentially including ECM-integrin and TNF superfamily receptor-ligand interactions. Interestingly, Treg support remains unique to the traditional immunomodulatory AFs, as profibrotic (TGFβ-treated) AFs lost their Treg-supportive capacity. This is also reflected in vivo, where inducible loss of fibroblast TGFβ signaling TGFbr2 cKO mice) led to enhanced immunomodulatory AFs and increased T2-Treg activation, ultimately improving lung health.

While we have demonstrated TGFβ can impact AF support of T2-Tregs, the cellular source and activators of TGFβ were not defined. Myeloid cells, such as macrophages, as well as Tregs themselves, are traditional sources of TGFβ^73^; however, TGFβ is secreted as a latent complex consisting of active TGFβ and the latency-associated peptide (LAP). Binding of LAP to integrins, including lung αvβ6 and αvβ8^74–76^, allows TGFβ to bind to its cell surface receptors and induce signaling. Tregs play crucial functional roles by activating TGFβ via αvβ8^77,78^. Therefore, while AFs directly support Tregs, we speculate that TGFβ activation from Tregs themselves plays a significant role in the local cross-regulation of AF function, thereby more broadly impacting adventitial immune “hubs”. T2-Tregs are well recognized for contributing to improved tissue health, with published mechanisms including enhanced immunosuppression and tissue repair via amphiregulin and IL-10 production. Our results show that the inducible loss of T2-Tregs led to reduced lung health after influenza infection, further highlighting the broader importance of T2-Tregs. Collectively, our data illustrate the diversity of lung fibroblast states after viral pneumonia and the crucial role of AFs in modulating key immune cells, such as Tregs, to preserve lung health. Modulation of fibroblast states, including the key role of AFs and their immunomodulatory properties, represents a promising future therapeutic target.

## Methods

### Mice

To trace regulatory T cells (Tregs), we used Foxp3*^eGFPcreERT2^* mice (JAX: 016961) intercrossed with Rosa26*^tdRFP^* (JAX: 038164), which carry an allele encoding a robust tandem-dimer RFP. These Foxp3*^creERT2^*; Rosa26*^RFP^* mice show RFP expression in *Foxp3*-lineage cells upon tamoxifen administration. Foxp3*^creERT2^*; Rosa26*^RFP^* mice were further crossed with an Il1r1*^eGFP^* mouse line (generated by Anna Molofsky lab, UCSF), a Rorc*^eGFP^*BAC transgene knock-in mouse line (MGI: 3829387), and a Tbx21*^ZsGreen^*transgenic mouse line (generously provided by JinFang Zhu, Lab of Immune System Biology, NIH). For our co-culture experiments and Treg isolation, we used Foxp3*^eGFP^* reporter mice (JAX: 006772). Foxp3*^creERT2^*; Rosa26*^RFP^* mice were crossed with Gata3*^flox^* mice (JAX: 028103) to specifically delete *Gata3* expression from *Foxp3*-lineage cells upon tamoxifen administration. Foxp3*^creERT2^*; Rosa26*^RFP^*mice were crossed to a PDGFRa^GFP^ knock-in knock-out mouse line (JAX: 007669), and used as heterozygotes, to visualize relative localization of Tregs to fibroblasts. To lineage trade sensory neurons, Nav1.8^cre^; tdTomato^f/+^ were used (generously provided by Sebastien Talbot, Department of Biological Sciences, Queens University).

To conditionally delete fibroblast *Tgfbr2*, Col1a2creERT2 mice were intercrossed with Tgfbr2^f/f^ mice (both Tgfbr2-exon2flox, MGI 2384513, and Tgfbr2-exon4flox, Jackson 012603). To conditionally delete adventitial fibroblasts, Gli1*^creERT2^* (JAX: 007913) mice were crossed with Rosa26*^DTR^* mice (JAX: 007900). To lineage trace profibrotic fibroblasts, we crossed Cthrc1*^CreER^*(generously provided by Dean Sheppard lab, UCSF) to Rosa26*^CAG-RFP^* (JAX: 007914). For the above conditional knockout strains, controls were littermate Cre-negative or flox-heterozygous mice.

The Saa3*^creER^* mouse strain was generously provided by Tatsuya Tukui (Dean Sheppard lab, UCSF) and generated by homology-directed repair at the endogenous Saa3 locus using CRISPR–Cas9 endonuclease activity in C57BL/6 mice as described previously^18^. In brief, CRISPR RNA (crRNA) with input sequence GTAGTTGCTCCTCTTCTCGG and trans-activating CRISPR RNA (tracrRNA) were obtained from IDT. A 5′ homology arm was amplified from C57BL/6 mouse genomic DNA with forward primer 5′-AGGTGAAATCTGTAGGTCAAAGATG-3′ and reverse primer 5′-GTATCTTTTAGGCAGGCCAGCAGGT-3′. A 3′ homology arm was amplified with forward primer 5′-GTTGTTCCCAGTCATGCTGC*TTC* CCGAGAAGAGGAGCAACTACTGGGT-3′ (*protospacer adjacent motif (PAM) sequence CCC was altered to TTC) and reverse primer 5′- GCAAAATTAGGGACCAATGACCTACTC-3′. A targeting vector with P2A-creERT2 sequence flanked by 5′ and 3′ homology arms was generated using NEBuilder

HiFi DNA Assembly (NEB) and cloned into a pKO2 backbone plasmid. The targeting vector was linearized at SalI (NEB) and NotI (NEB) sites flanking the donor DNA sequence and the linearized donor DNA was purified by agarose gel electrophoresis with GeneJet Gel Extraction kit (Thermo Fisher). Linearized donor DNA and CRISPR–Cas9 complex were injected into C57BL/6 fertilized zygotes, which were then implanted into the oviducts of pseudopregnant female mice. A total of 195 embryos were implanted and 15 pups were born. Two founders were identified by genotyping. We used one founder to expand the colony.

Mice were mixed-sex, backcrossed onto C57BL/6 for at least 10 generations. If not otherwise stated, all experiments were performed with 8–16-week-old male and female mice. All mice were bred and maintained in specific-pathogen-free conditions at the animal facilities of UCSF and were used in accordance with institutional guidelines and under study protocols approved by the UCSF Institutional Animal Care and Use Committee.

### Tamoxifen-induced Cre recombination

Mice were injected intraperitoneal (i.p.) with 200mL of tamoxifen (Sigma-Aldrich) dissolved in corn oil at 10mg/mL.

### Influenza A (PR8) infection

Influenza A/Puerto Rico 8/1934 (PR8) H1N1 strain was obtained from ATCC (VR-95), aliquoted and stored in liquid nitrogen. Mice were anesthetized by i.p. injection with ketamine–xylazine (Patterson Veterinary) or isoflurane and intranasally infected with ∼60 plaque-forming units for each experiment. Mice were sacrificed at the indicated time points and tissues were harvested and analyzed.

### qPCR analysis

Lung lobes taken for qPCR were snap-frozen at -20°C in 1.5mL Eppendorf tubes. Lobes were then homogenized in 600mL of buffer RLT (RNeasy Mini Kit, Benchmark D1032-30) with a Precellys Evolution Touch Homogenizer (Bertin Technologies). Lysates were then used to extract RNA according to the manufacturer’s protocol of the RNeasy Mini Kit. Eluted RNA was then converted to cDNA using SuperScript III First Strand Synthesis System (Invitrogen). qPCR was run on a StepOnePlus Real-Time PCR system (Applied Biosystems).

Primers used: NP-1 (5’-CAGCCTAATCAGACCAAATG-3’ (forward) and 5’-TACCTGCTTCTCAGTTCAAG-3’ (reverse)),

### Tissue processing: Thick section imaging

Following CO2 euthanasia, mice were transcardially perfused with 10mL of DPBS and, if not used for flow cytometry, flushed intra-tracheally with 4% PFA (Thermo Scientific). Lung lobes were then stored in 4% PFA for 1-3 days at 4°C followed by cryoprotection (30% sucrose). Lobes were frozen in O.C.T. (Thermo Scientific) on dry ice and sliced into 300mM sections on a cryostat (Leica). Sections were then placed into PBS to wash and blocked (DPBS, 1% BSA, 0.3% TritonX-100, 1% mouse serum) overnight at 4°C. Sections were then incubated in the primary antibody diluted in blocking buffer for 3 days at 4°C. Sections are then washed with washing buffer (DPBS, 0.3% TritonX-100, 0.5% 1-Thioglycerol) for three 30-minute washes at 37°C. Sections were then incubated overnight at 4 °C with a secondary antibody solution diluted in blocking buffer. Sections were washed once again for three 30-minute washes and then cleared using Ce3D clearing solution (for 5mL: 2.75mL of 40% N-methylacetamide, 80% weight/volume Histodenz, 5mL of TritonX-100, 25mL of thioglycerol) for one hour at room temperature. Sections were then mounted and imaged with a Leica SP8 confocal microscope.

### Imaging Antibodies

Primary antibodies used for murine imaging include chicken anti-GFP (Aves Labs GFP-1020, 1:200), rabbit anti-dsRed (Takara 632496, 1:300), rat anti-ER-TR7 (Novus Biologicals NG100-64932), goat anti-Pi16 (R&D Systems AF4929), rabbit anti-proSP-C (Millipore AB3786), goat anti-IL-33 (R&D Systems AF3626), and rat anti-EpCAM (BD Pharmingen 552370).

Directly conjugated antibodies were also used including mouse anti-αSMA AF488 (eBioscience 53-9760-82), anti-αSMA AF405 (R&D Systems IC1420V025), rat anti-CD45 BV421 (BioLegend 103134), rat anti Lyve1 ef660 (eBioscience 50-0443-82) and rat anti-CD45 (BioLegend 103124).

Secondary antibodies were used as necessary at protocol-specific concentrations, conjugated to AF488, AF555, AF594, and AF647 (Life Technologies, Thermo-Fisher).

### Confocal microscopy

Confocal images (for thick sections and quantification) were acquired using a Leica SP8 laser scanning confocal microscope equipped with an AOBS tunable detection pathway, a white-light laser that excites between 470-670 nanometers, and a lighting deconvolution module. An Apo IMM (H2O Dipping) 20x 1.95mm working distance objective was used. Z steps were acquired every 2.5 or 5uM.

### Image Analysis/Quantification

Z-stacks were rendered in 3D and quantitatively analyzed using Bitplane Imaris v9.8 software package (Andor Technology OLC, Belfast, N. Ireland). Individual cells (e.g., lymphocytes) were annotated using the Imaris surface function based on fluorescent reporter signal (e.g., Rosa26^RFP^) when available, along with DAPI staining, and size/morphological characteristics, with background signal in unrelated channels excluded. 3D reconstructions of vessels and airways were generated using the Imaris surface function based on fluorescent αSMA signal. 3D distances between Tregs and αSMA surfaces were calculated using the Imaris Distance Transform Matlab extension. Treg quantifications and distance calculations were performed across various lung slices, prioritizing regions balanced between adventitial and alveolar regions. Similar surfacing thresholds were applied across slices, and experiments for given cell types and structures, and individual thresholds were adjusted when necessary to account for varying tissue background. Damage patches were quantified by manually drawing surfaces based on a combination of pro-SPC and CD45 fluorescent signal. Another surface would then be drawn for the full lung slice, and a percentage was calculated by dividing the volume of the damage patch by the volume of the whole lung.

### Tissue processing: single cell isolation (murine)

Single-cell tissues were prepared from tissues including lung, spleen, and draining lymph nodes (dLN). Immediately following CO2 euthanasia, spleens were removed and put into 1X DPBS. Mice were subsequently transcardially perfused through the left ventricle with 10mL of DPBS. Lungs were then removed, as well as the mediastinal lymph nodes, and put into DPBS. Spleen and lymph nodes were crushed through 70mM filters using 3mL syringes and washed with FACS buffer (3% FBS, 0.5% NaN3). Lungs were put into gentleMACS tubes (Miltenyi Biotec 130-096-334) with 5mL of digestion media (40mg/mL LiberaseTM and 80mg/mL of DNase I (Roche) in HBSS). Tubes were then placed on a gentleMACS Octo Dissociator (Miltenyi Biotec 130-096-427) and spun. Tubes were then placed at 37°C for 30 minutes and spun once more on the gentleMACS Octo Dissociator. Digested tissue was subsequently filtered through 70 μm filters using 3mL syringes and washed with FACS buffer. After filtration, samples are centrifuged at 1500rpm for 5 minutes at 4°C and resuspended in 1mL of 1X Pharmlyse buffer (BD Biosciences 555899) for 2 minutes at 4°C (spleen and dLN skip the Pharmlyse step). Samples are then quenched with 6mL of FACS buffer and centrifuged once more at 1500rpm for 5 minutes at 4°C. Samples are then resuspended to the necessary concentration for staining.

### Flow cytometry

Resuspended samples were stained in 96-well V-bottom plates. Surface staining was performed at 4°C for 45 minutes in 50mL staining volume. For experiments involving intracellular staining, cells were fixed and permeabilized using Foxp3 Transcription Factor Staining Buffer Set (eBioscience) followed by staining at 4°C for 1 hour or overnight in 50mL staining volume. All samples were acquired on a BD LSRII Fortessa Dual or a BD FACSAria II for cell sorting. Live cells were gated based on their forward and side scatter followed by Zombie NIR fixable (BioLegend 423106), Fixable Viability Dye eF780 (eBioScience 65086514), Draq7 (Biolegend 424991), or DAPI (40,6-diamidine-20-phenylindole dihydrochloride; Millipore Sigma D9542-10MG) exclusion. Lineages were subsequently identified as follows:

T cells were identified as CD45+, CD11b-, CD19-, NK1.1-, CD3e+, CD4+ (CD4 T cells), CD8a+ (CD8 T cells), Foxp3^+^ (Tregs), and were further defined as CD44^-^CD62L+ (naïve T cells), CD62L-, CD44+ (activated T cells), or CTV diluted (proliferating T cells). Additionally, CD4 T cells and Tregs were defined as T-bet+ (Th1 T cells), Gata3+ or ST2^+^ (Th2 T cells), or RORgt+ (Th17 T cells). Neutrophils were defined as CD45+, CD11b+, Ly6G+ (and optionally Thy1-, CD19-, NK1.1-). Monocytes were defined as CD45+, CD11b+, Ly6G-, Ly6C+ (and optionally Thy1-, CD19-, NK1.1-, Siglec F-). Macrophages were defined as CD45+, Ly6G-, Ly6C-, CD64+ (optionally MERTK+). cDCs were identified as CD45+, Ly6G-, Ly6C-, CD64-, MHCII+, CD11c+, and were further defined as cDC1s (CD11b^lo^, optionally SIRPa-) or cDC2s (CD11b^hi^, optionally SIRPa+). B cells were defined as CD45+, Thy1-, CD19+. Eosinophils were defined as CD45+, Thy1-, CD19-, NK1.1-, Ly6G-, CD11b+, Siglec F+. Populations were backgated to verify purity and gating.

Data were analyzed using FlowJo (Treestar, USA) and compiled in Prism (GraphPad) software. Cell counts were performed using flow cytometry counting beads (CountBright Absolute; Life Technologies) according to the manufacturer’s instructions.

### Flow Cytometry Antibodies

Antibodies used for flow cytometry include anti-CD45 (30-F11, BD Biosciences 564279), anti-CD90.2 (Thy1, 53-2.1, Biolegend 140327, BD Biosciences 553004), anti-CD11b (M1/70, Biolegend 101224 or BD Biosciences 563015), anti-CD19 (6D5, Biolegend 115554), anti-NK1.1 (PK136, Biolegend 108736), anti-CD3e (17A2, Biolegend 100216), anti-CD4 (RM4-5, Biolegend 100557, or GK1.5, BD Biosciences 563050), anti-CD8a (53-6.7, Biolegend 100750), anti-CD44 (IM7, Biolegend 103030), anti-CD69 (H1.2F3, Biolegend 104505), anti-CD62L (MEL-14, Biolegend 104407), anti-Tbet (4B10, Biolegend 25-5825-80), anti-Gata3 (TWAJ, eBioscience 12- 9966-41), anti-RORgt (B2D, eBioscience 17-6981-82), anti-Ly6G (1A8, Biolegend 127624), anti-Ly6C (HK1.4, Biolegend 128011 or 128035), anti-CD64 (X-54-5/7.1, Biolegend 139323 or BD Biosciences 558539), anti-MERTK (DS5MMER, eBioscience 46-5751-80), anti CD11c (N418, Biolegend 117339 or 117318), anti-CD172a (SIRPa, P84, eBioscience 12-1721-80), anti-Siglec-F (E50-2440, BD Biosciences 740956), anti-I-A/I-E (MHCII, M5/114.15.2, BD Biosciences 748845), anti-CD192 (CCR2, 475301, BD Biosciences 747964), anti-Foxp3 (FJK-16S, Invitrogen 53-5773-82), anti-gdTCR (GL3, Invitrogen 14-5711-82), anti-CD25 (PC61, BioLegend 102030), anti-ST2 (DJ8, MD BioProducts 101001PE), anti-CD152 (CTLA-4, UC10-4B9, BioLegend 106310), anti-KLRG1 (MAFA, BioLegend 138421), anit-CD278 (ICOS, C39A.4A, BioLegend 313537), anti-CD279 (PD-1, 29F.1A12, BioLegend 135225), anti-Ki67 (B56, BD Biosciences 564071), anti-podoplanin (gp38, 8.1.1, BioLegend 127411), anti-PDGFRa (APA5, BioLegend 135908), anti-CD326 (EpCAM, G8.8 BioLegend 118320), biotinylated anti-CD31 (390, BioLegend 102404), biotinylated anti-CD45 (30-F11, BioLegend 103104), and anti-Ly-6A (Sca-1, D7, BioLegend 108131).

### Ex vivo co-culture

For *ex vivo* co-culture experiments, lung adventitial fibroblasts (AFs) were obtained by dissection as described above but using a different digest media. The digest media per mouse was instead: 5mL 1X Dispase II, 112.5 U/mL Collagenase I, and 40ug/mL DNase I. After cardiac perfusion, 1.5mL of digest media was injected through the trachea to expand the lungs. Lungs were then chopped manually with sterile razors to small chunks and placed into digestion media at 37°C for 30 minutes. Samples were then pushed through a sterile 70mM filter with a 3mL syringe and centrifuged at 1500rpm at 4°C for 5 minutes. Samples were resuspended in sterile Pharmlyse, placed on ice for 1 minute, then quenched and centrifuged again. Samples were resuspended in sterile FACS buffer at 1mL/lung added and incubated with 4mL/Lung of rat serum (Thermo Fisher 10710C) on ice for 10 minutes. Biotinylated anti-CD45 and anti-CD31 are added at 4mL/mL of cells and incubated for 30 minutes on ice. Biotinylated cells were subsequently removed by using EasySep Magnetic Beads (Stem Cell), according to the manufacturer’s instructions. AFs were plated at 10,000 cells/well in a flat-bottom 96-well plate in AF media (DMEM, 10% FBS, 1X Glutamax, and 1% penicillin/streptomycin) and incubated at 37°C, 5% CO2 for 7 days, where regulatory T cells were added.

To sort Tregs, T cells were isolated from the spleen and cervical/inguinal lymph nodes of reporter mice using EasySep Magnetic Bead negative selection, and the cells were subsequently stained for sorting. Tregs were labeled with 5mM CellTrace Violet in PBS (Thermo Scientific) for 10 minutes, followed by washing with Treg media (RPMI, 10% FBS, 1X Glutamax, 1% penicillin/streptomycin, 25mM HEPES, and 1X NEAA), centrifugation, and resuspension. Tregs were plated at 10,000 cells/well in all conditions. For all Treg wells, 1000IU of IL-2 was added to ensure healthy Treg populations. As a positive control, anti-CD3/CD28 T-cell activating DynaBeads (Thermo Scientific) were magnetically washed in 1mL DPBS and added at 3:1 bead:cell ratio. After 72 hours of co-culture, wells were mixed by pipetting, and cells were transferred to a V-bottom plate. The remaining cells were trypsinized for 10 minutes at 37°C, quenched, centrifuged, and added to the previous cells. Cells were then centrifuged and stained for flow cytometry.

For AFs treated with TGFβ, on Day 3 post-plating, 100mL of media was removed and 100mL of media containing the relevant amount of TGFβ was added. On the day of Treg addition, the TGFβ-treated media was completely removed, and wells were washed with PBS before addition of Tregs. For IL-33-treated Tregs, IL-33 was added at the same time as IL-2 into the Treg media in its relative amounts.

### Murine scRNAseq data processing

Sequencing data were aligned to mouse genome mm10 with CellRanger version 7.1.0 (scRNAseq experiment 1; time course of global cells) or version 7.2.0 (scRNAseq experiment 2; time course of stromal cells) (10x Genomics). Data was processed using the Seurat R package, version 4.3.3 (experiment 1) or 5.1.1 (experiment 2). We excluded cells with high mitochondrial gene expression and low or high UMI and feature counts, using the bottom and top 2.5 percentiles as our cutoff. *SCTransform* function was performed before clustering. Principle components were calculated using Seurat’s *RunPCA* function, followed by graph-based clustering using Seurat’s *FindNeighbors* (dims =1:30 for experiment 1, dims = 1:15 for experiment 2) and *FindClusters* (res = 0.25 for experiment 1, res = 0.8 for experiment 2) functions and 2D visualization using Seurat’s *RunUMAP* function (dims = 1:30 for experiment 1, dims = 1:15 for experiment 2). We used a combination of *FeaturePlot*, *DotPlot,* and *VlnPlot* to identify cell clusters (for both experiments) using common and well-validated lineage marker genes (e.g., *Col1a1, Col1a2, Pdgfra* [fibroblasts]; *Pecam1* [endothelial]; *Itgam, Itgax* [myeloid]; *Cd3e, Cd4, Cd8* [T cells]; *Sftpc* [epithelial]; *Scube2, Ces1d* [alveolar fibroblasts]; *Pi16, Ccl11, Col14a1* [adventitial fibroblasts]; *Saa3, Ccl2, Ifitm3* [inflammatory fibroblasts]; *Pdgfrb, Cox4i2* [pericytes]; *Acta2* [smooth muscle]; *Acta2, Hhip* [peribronchial], abd *Msln, Wt1* [mesothelial]). We identified and excluded 2 clusters made up of likely doublets and contaminants based on gene expression, visualizing resultant clusters on UMAPs and dot plots using Seurat. Final datasets comprised 31,603 cells including 19,381 lymphocytes; 7472 myeloid cells; 2862 fibroblasts; 1712 endothelial cells; and 176 epithelial cells (experiment 1) and 34,995 cells including 18,793 alveolar fibroblasts; 11,886 adventitial fibroblasts; 1362 inflammatory fibroblasts; 1138 pericytes; 760 smooth muscle cells; 553 peribronchial cells; 392 eptihelial cells; and 111 mesothelial cells (experiment 2). Additional feature plots, UMAP plots and dot plots, and heatmaps were generated using Seurat.

Fibroblasts and immune cells were subset and reclustered as above. In experiment 1, Tregs were found by reclustering the lymphocyte population (dims = 1:15, res = 0.8) and a combination of *FeaturePlot*, *DotPlot,* and *VlnPlot* to identify clusters high for *Foxp3* expression. Clusters were isolated and reclustered again (dims = 1:10, res = 0.9) and a combination of *FeaturePlot*, *DotPlot,* and *VlnPlot* to identify clusters high for *Foxp3* expression was once again used and isolated for final downstream analysis. Fibroblasts were found by reclustering the fibroblast population (dims = 1:10, res = 0.8) and a combination of *FeaturePlot*, *DotPlot,* and *VlnPlot* to identify clusters high for canonical fibroblast markers. Clusters were isolated and reclustered again (1:10, res = 0.8) for downstream analysis. In experiment 2, adventitial fibroblasts were isolated and reclustered (dims = 1:15, res = 0.8) for downstream analysis. Clusters expressing high levels of myeloid or genes were excluded as likely contaminants and removed from global UMAP and bar plots. For experiment 2, in the adventitial fibroblast cluster, clusters of different AF states were found by using *AddModuleScore* (selecting for genes unique to each AF state such *Aqp1* and *Il33*) and violin plots. Relative abundance across time was calculated for each fibroblast subcluster after normalizing for timepoint or condition sample size (i.e., total # cells). Multi-gene scores were calculated as follows and generated using *AddModuleScore.* Profibrotic fibroblasts scores were calculated using a combination of the following genes: *Cthrc1, Acta2, Col1a1, Vim, Postn, Tnc, Lrrc15,* and *Spp1*. Type 2-like Treg scores were calculated using a combination of the following genes which are known to associate with type 2 immunity or type 2-like Tregs: *Gata3, Il1rl1, Areg, Prdm1, Il2ra, Irf4,* and *Setd2*^71,79^. Similarly, type 1-like Treg scores were calculated using a combination of the following genes: *Tbx21, Cxcr3, Ifng, Cx3cr1, Il10,* and *Tgfb1*. Finally, AF immune-functionality scores were calculated using a combination of markers seen to be unique to immune-modulatory AFs: *Il33, Ptgs2, Lum,* and *Ccl11*.

### Xenium spatial transcriptomics

A custom-designed 480 gene stromal-immune panel (Table somewhere) was generated using insights from single cell studies (see sc refs in paper, Anthony’s and Dean Sheppard’s). Following CO2 euthanasia, mice were transcardially perfused with 10mL of D-PBS and lung lobes were collected. Tissues were immediately fixed in 4% PFA and left to fix for 24h at 4C. Samples were washed 3x times with 1x D-PBS before transfer to 70% Ethanol and FFPE blocks were made (using UCSF Gladstone histology core, need to cite). Once blocks were generated, we partnered with UCSF Genomics CoLabs(citation) to process and analyze samples. In brief, FFPE blocks were sectioned using a microtome at 10-μm thickness, placed on Xenium slides, deparaffinized and de-crosslinked according to manufacturer instructions (CG000578, Rev F, 10x Genomics). Probes were hybridized, ligated, and amplified on slide, cell segmentation staining and amplification quenching were performed (CG000749, Rev B, 10x Genomics) and slides were analyzed with a Xenium Analyzer (CG000584, 10X Genomics). After scanning, slides were removed from the analyzer and subjected to post-run H&E per 10x Genomics protocol (CG000613, Rev B, 10x Genomics).

Raw Xenium Data was loaded into RStudio as a Seurat object using *LoadXenium* function. Cells with <10 counts, <5 features or the top 2% of outliers in either metric were excluded per slide. Following exclusion Seurat objects were merged into a single dataset and log-normalized using Seurat’s *NormalizeData* function, variable features were identified, and *ScaleData* function was applied. Principle components were calculated using Seurat’s *RunPCA* function, followed by graph-based clustering with Seurat’s *FindNeighbors* (dims 1:20 k.param = 16) and *FindClusters*(res = 0.5) functions. Finally, data was visualized in 2D with Seurat’s RunUMAP function (dims 1:20) and *JoinLayers* was run. To aid in cell type identification, we used *FindAllMarkers* to identify cluster-defining DEGs. Further cell-identification workflow follows as described in the scRNAseq data processing method. Clusters were spatially visualized using the 10x Xenium Explorer tool.

## Acknowledgments

We thank Dr. Richard Locksley for his constructive feedback on this manuscript, as well as the UCSF Parnassus Flow Core RRID:SCR018206 and DRC Center Grant NIH P30 DK063720, the UCSF Biological Imaging and Development Core (BIDC), the UCSF Genomics CoLab, the UCSF Institute for Human Genetics, and Histowiz, Inc. for instruments and services. D.S. is supported by the NHLBI (NIH R01HL142568). A.B.M., and A.V.M. are supported by the NINDS (NIH R01NS126765), UCSF PBBR grant, and UCSF Dept. of Lab Medicine discretionary funds; A.B.M. is additionally supported by the NIAID (NIH R01AI162806).

## Statistical analysis

All data were analyzed by comparison of means using unpaired (unless otherwise noted) two-tailed Student’s t-tests; for multiple comparisons, one-way or two-way ANOVA (with Tukey or Sidak’s post hoc test) was used as appropriate (Prism, GraphPad Software, La Jolla, CA). ns = not significant, *p<0.05, **p<0.01, ***p<0.001, ****p<0.0001. Figures display means ± standard deviation (SD) unless otherwise noted. When possible, results from independent experiments were pooled. All data points reflect individual biological mouse replicates (flow analysis) or individual tissue slices analyzed (imaging), unless otherwise noted. For imaging experiments, all analysis was performed with at least three mice from at least two independent experiments, with two or more fields analyzed per mouse.

---

**Figure S1:**
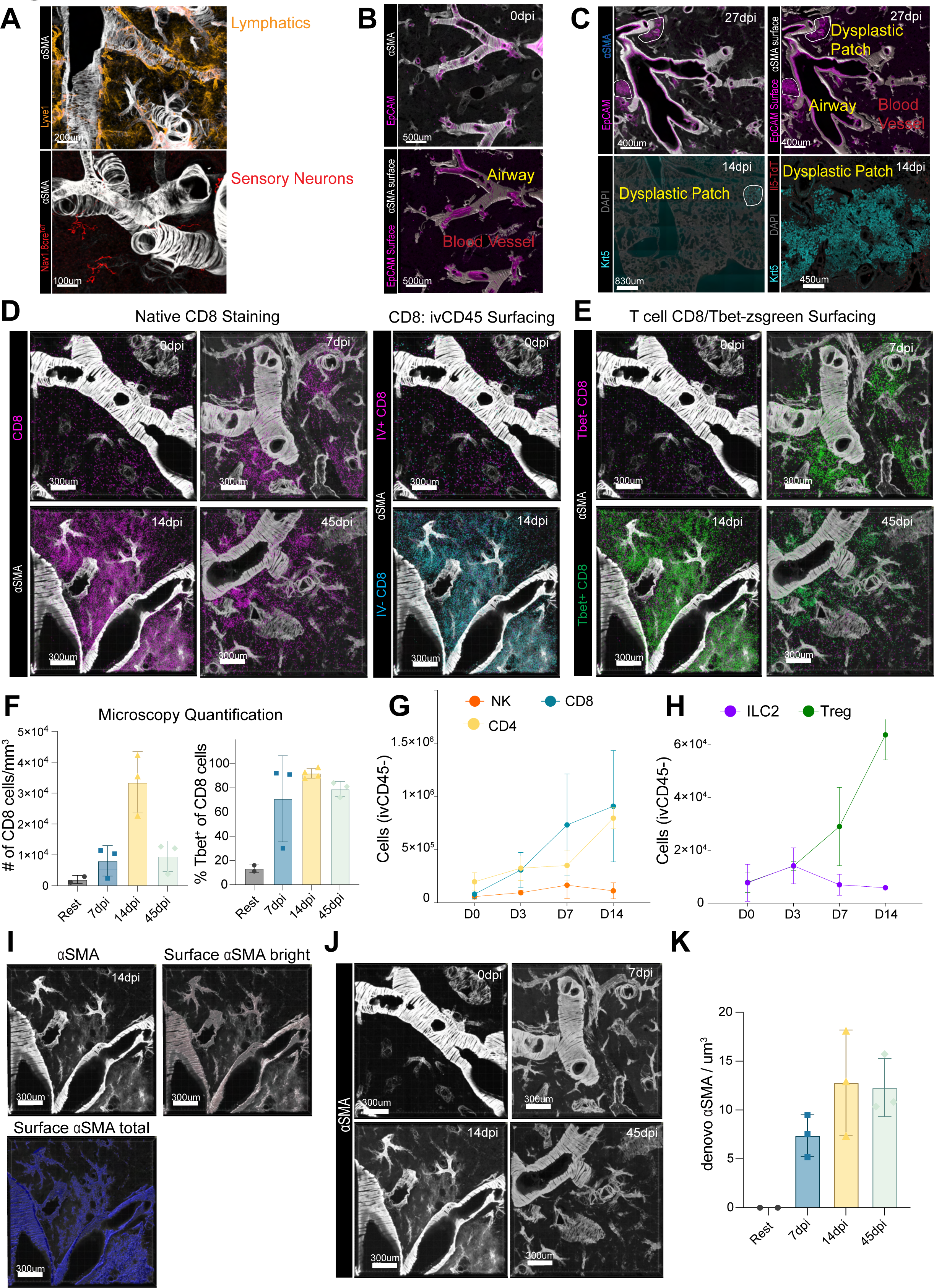
**A**, Thick section imaging illustrating localization of lymphatics (Lyve1^+^) and sensory neurons (Nav1.8^+^). **B**, Thick section imaging and Imaris surfaces illustrating blood (αSMA^+^EpCAM^-^) and airway (αSMA^+^EpCAM^+^) vasculature at 0 dpi. **C**, Thick section imaging and Imaris surfaces illustrating the development of dysplastic patches at both 14 and 27 dpi. **D**, Thick section imaging of expansion and contraction of CD8 T cells (CD8^cre^; Rosa26^tdt^; Tbet^ZsGreen^ reporter mice) across timepoints after influenza infection (left panels), and of expansion of IV^-^ CD8 T cells between 0 and 14 dpi (right panels). **E**, Thick section imaging illustrating expansion and contraction of Tbet^+^ (green) vs. Tbet^-^(magenta) CD8 T cells across timepoints after influenza infection. **F**, Quantitation from thick section imaging illustrating standardized numbers and frequencies of CD8 cells across influenza time course. **G-H**, Flow cytometry data measuring numbers of (**G**) NK, CD8, and CD4 (CD4^+^Foxp3^-^) and (**H**) Tregs, and ILC2s across timepoints after influenza infection. **I**, Thick section imaging identifying de novo αSMA generation (αSMA^dim^) versus traditional vasculature (αSMA^bright^) at 14 dpi. **J**, native staining of αSMA in thick section imaging across influenza infection. **K**, Quantification of de novo αSMA surfacing across influenza infection. **A-E**, and **I-J**, 200-300µM slices; images represent 2 or more mice.

**Figure S2:**
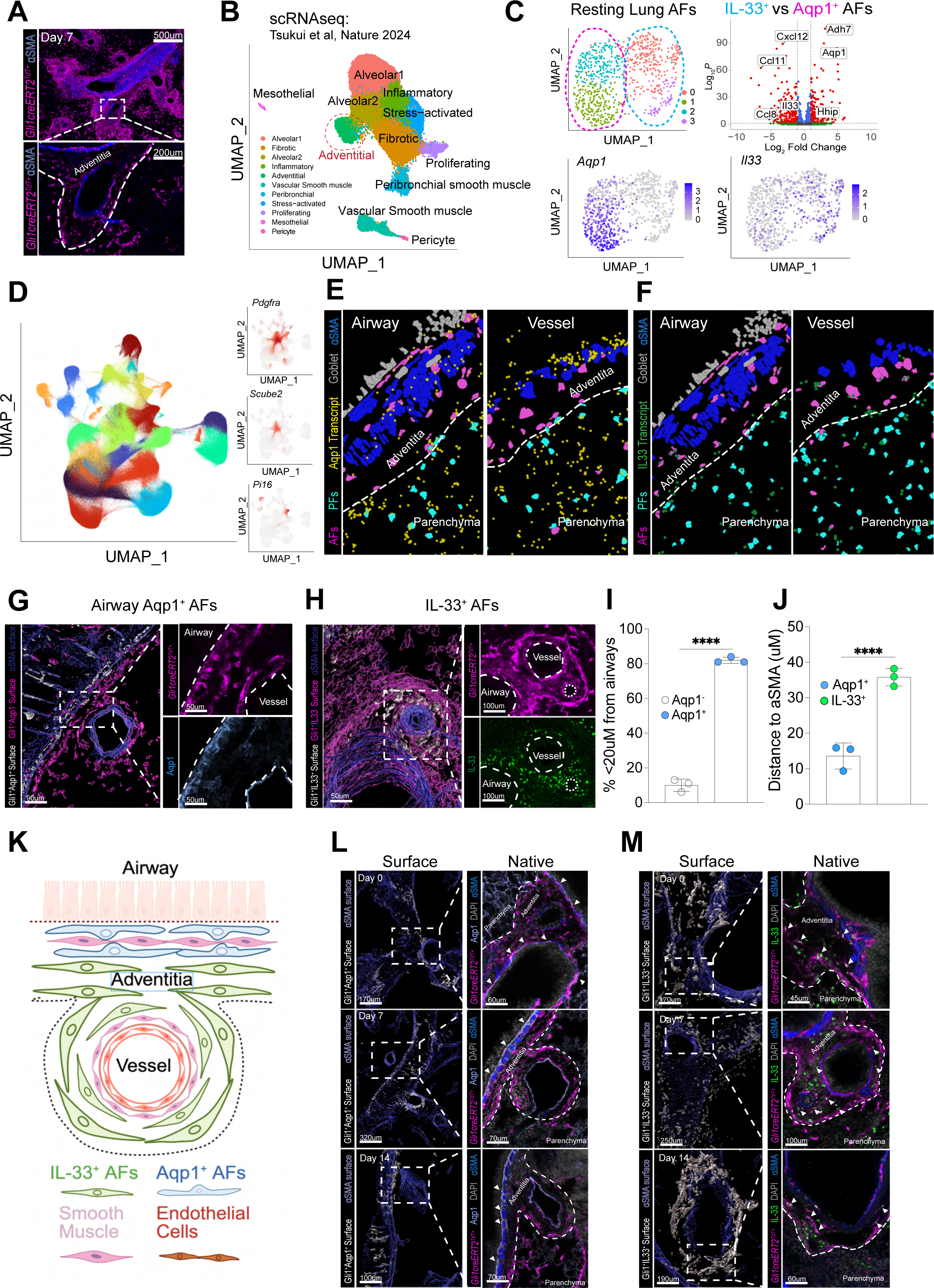
A, Thick section and zoomed image of adventitial spaces marked by Gli1 lineage tracing (white dashed line). **B**, UMAP plot (Tatusya et al., 2024) demarcating various cell populations sorted from healthy and bleomycin-treated mice. **C**, UMAP plot and volcano plot illustrating transcriptional differences between IL-33^+^ and Aqp1^+^ AFs. **D**, UMAP plot of total Xenium spatial sequencing data illustrating the location of clusters of alveolar (Scube2^+^) and adventitial (Pi16^+^) fibroblasts. **E**, Magnified spatial sequencing image illustrating differences of *Aqp1* transcript (yellow) overlap over AFs (magenta) between airway (left panel) and blood vessel (right panel) regions. **F**, Magnified spatial sequencing image illustrating differences of *Il33* (green) transcript overlap over AFs (magenta) between airway (left panel) and blood vessel (right panel) regions. **G**, Imaris surfaces looking at relative localizations of Aqp1^+^ (white) and Aqp1^-^ (magenta) AFs to αSMA (left panel) with native staining of Gli1 lineage tracer (magenta, top right) and Aqp1 (white, bottom right). **H**, Imaris surfaces showing relative localizations of IL-33^+^ (white) and IL-33^-^(magenta) AFs to αSMA (left panel), with native staining for the Gli1 lineage tracer (magenta, top right) and IL-33 (green, bottom right). **I**, Imaris quantitation illustrating frequency of Aqp1^+^ and Aqp1^-^ surfaces that are less than 20µM from identified airways (using DAPI method). **J**, Imaris quantitation illustrating average distances of Aqp1^+^ and IL-33^+^ AFs to αSMA. **K**, Graphical illustration showing differential localization of Aqp1^+^ and IL-33^+^ AFs within the adventitia. **L**, Time course series of images looking at 0, 7, and 14 dpi timepoints of airway-associated (Gli1^+^Aqp1^+^) AFs. Left panels illustrate Imaris surfaces showing αSMA surfaces (blue) and airway-associated AF surfaces (white). Right panels illustrate native immunofluorescence (IF) and reporter staining of overlapping Gli1 staining (magenta) and Aqp1 staining (sky blue) in adventitial regions (white dashed line). **M**, Time course series of images looking at 0, 7, and 14 dpi timepoints of immune-modulatory (Gli1^+^IL-33^+^) AFs. Left panels illustrate Imaris surfaces showing αSMA (blue) and immune-modulatory (white) surfaces. Right panels illustrate native immunofluorescence (IF) and reporter staining of overlapping Gli1 staining (magenta) and IL-33 staining (green) in adventitial regions (white dashed line). (**A, G-H, L-M**) 200-300µM slices. (**E-F**) 20-40µM slices; images represent two or more mice. (**I-J**) Graph shows mean±s.d. ****p<0.0001; paired Student’s T-test.

**Figure S3:**
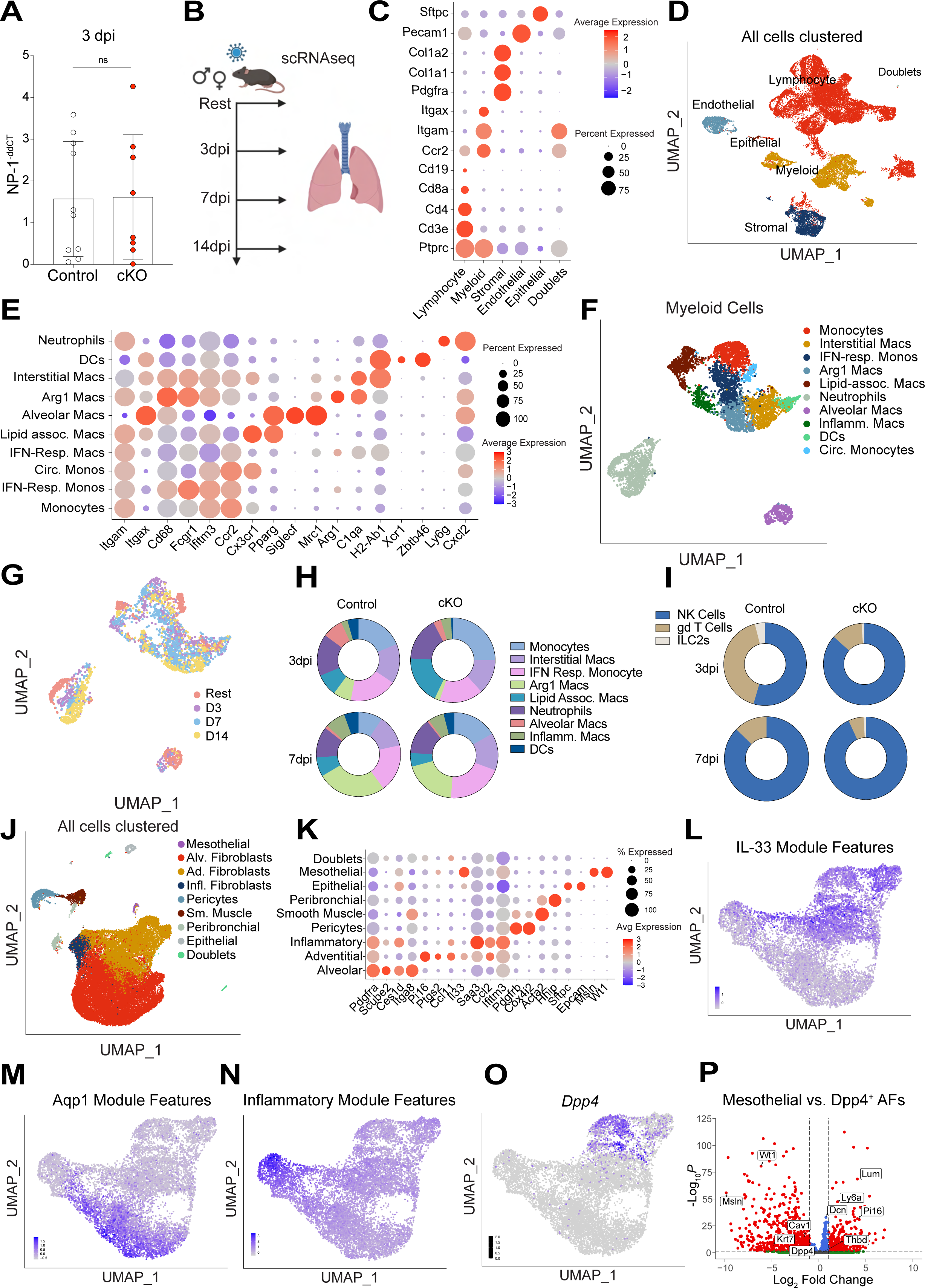
**A**, qPCR quantitation of 3 dpi viral levels using NP-1 primers between control and cKO mice. **B**, Graphical illustration of scRNAseq experiment. Sorting strategy: 5 populations of cells were sorted in equal numbers and combined for sequencing: CD4 T cells (CD3^+^CD4^+^), Total lymphocytes (CD45^+^Thy1.1^+^), Myeloid cells (CD45^+^Thy1.1^-^), Stromal cells (CD45^-^PDGFRα^+^) and other lineage negative structural cells (CD45^-^ PDGFRα^-^). All littermate mice were infected on the same day and harvested at 0, 3, 7, and 14 dpi. **C-D**, Dot plot (**C**) and UMAP (**D**) illustrating the various cell populations sorted out: Lymphocytes, myeloid, fibroblasts, endothelial, epithelial, doublets, and the marker genes for all cellular clusters. **E-F**, Dot plot (**E**) and UMAP (**F**) identifying various myeloid cell populations (**F**): Monocytes, interstitial macrophages, IFN-responsive monocytes, Arg1^+^ macrophages, lipid-associated macrophages, neutrophils, alveolar macrophages, inflammatory macrophages, dendritic cells, and circulatory monocytes, and their marker genes. **G**, UMAP of myeloid clusters illustrating timepoint metadata. **H**, Pie chart showing relative proportions of all identified myeloid cells comparing control to TGFbr2 cKO mice at 3 and 7 dpi. **I**, Pie chart showing relative proportions of all identified innate lymphoid cells (NK cells, ψ8 T cells, and ILC2s) comparing control to TGFbr2 cKO mice at 3 and 7 dpi. **J-K**, Global UMAP (**J**) and (**K**) Dot plot illustrating the various cell populations sorted out: mesothelial cells, alveolar fibroblasts, adventitial fibroblasts, inflammatory fibroblasts, pericytes, smooth muscle cells, peribronchial (sub-epithelial) fibroblasts, epithelial cells, and doublets, and their marker genes. **L-O**, UMAP representation of (**L**) IL-33 immune-modulatory module score, (**M**) *Dpp4* gene expression, (**N**) Aqp1^+^ airway-associated module score, and (**O**) inflammatory module score. **P**, Volcano plot comparing DEGs between traditional mesothelial cluster and fascial-like Dpp4^+^ AF cluster. ns = not significant; Student’s T-test (**A**).

**Figure S4:**
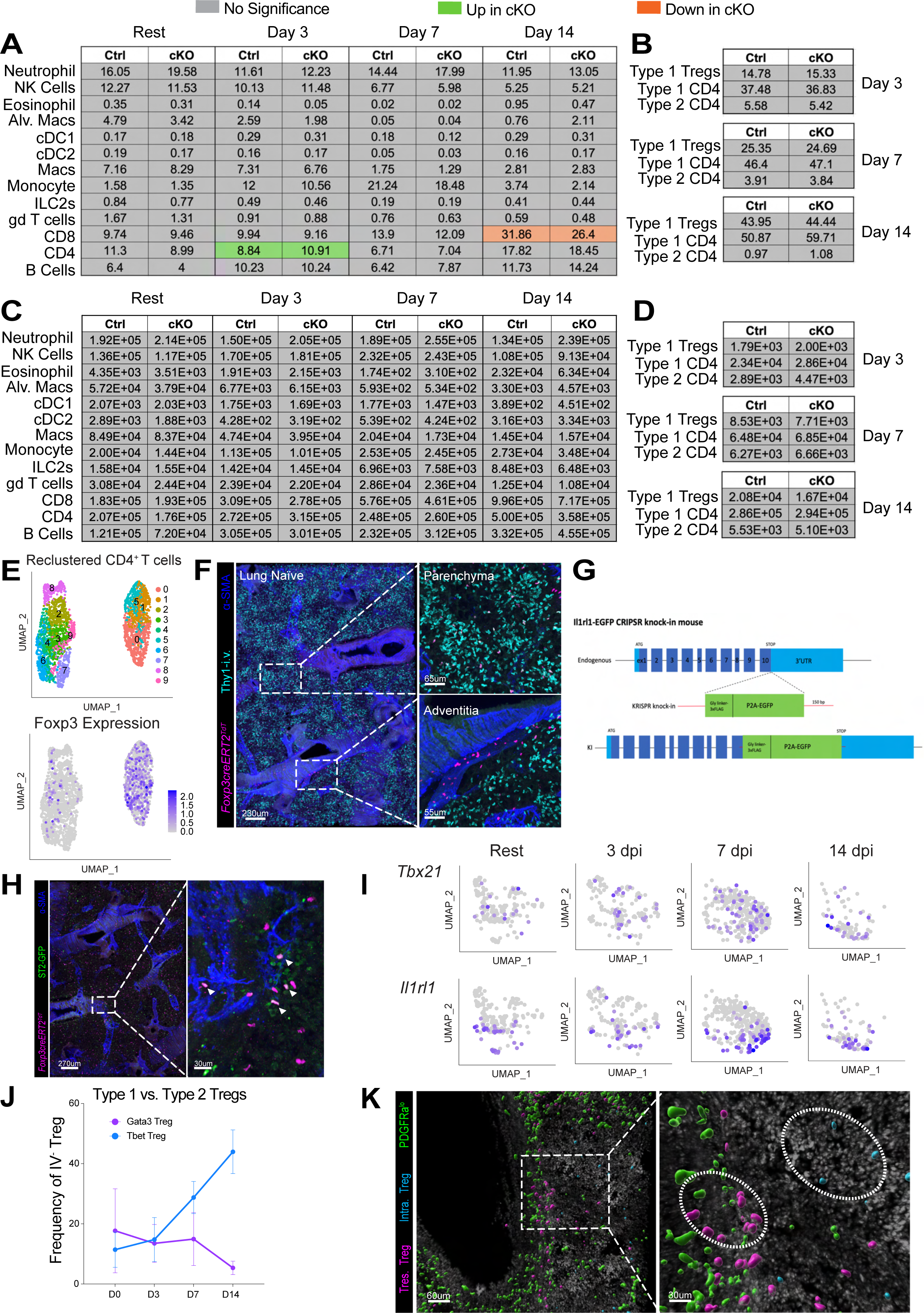
**A-D**, Tables illustrating values of time course flow cytometry data looking at frequencies (**A-B**) and cell number (**C-D**) of various immune cells between control and TGFbr2 cKO mice at rest, 3, 7, and 14 dpi. **E**, UMAP illustrating filtered and re-clustered CD4^+^ cells and *Foxp3* expression for Tregs. **F**, Representative imaging of CD45/Thy1.2i.v. labeling illustrating intravascular Tregs (i.v.^+^, top right) and tissue resident Tregs (i.v.^-^, bottom right). **G**, Graphical illustration of ST2^GFP^ reporter construct. **H**, Thick-section native imaging illustrating type 2 Tregs (ST2^GFP+^Foxp3^CreERT2^; Rosa26^RFP+^, white arrows). **I**, Serial UMAPS illustrating *Tbx21* and *Il1rl1* expression in Tregs over time after influenza. **J**, FACS data illustrating frequencies of Type 1 (blue) and Type 2 (purple) Tregs at rest, 3, 7 and 14 days post influenza infection. **K**, Representative image showing Imaris surfacing of PDGFRα^lo^ cells (green) and Tregs (magenta). Graphs show mean±s.d.; Student’s T-test for each time point (**A-D**). (**F, H, I**) 200-300µM slices; images represent two or more mice.

**Figure S5:**
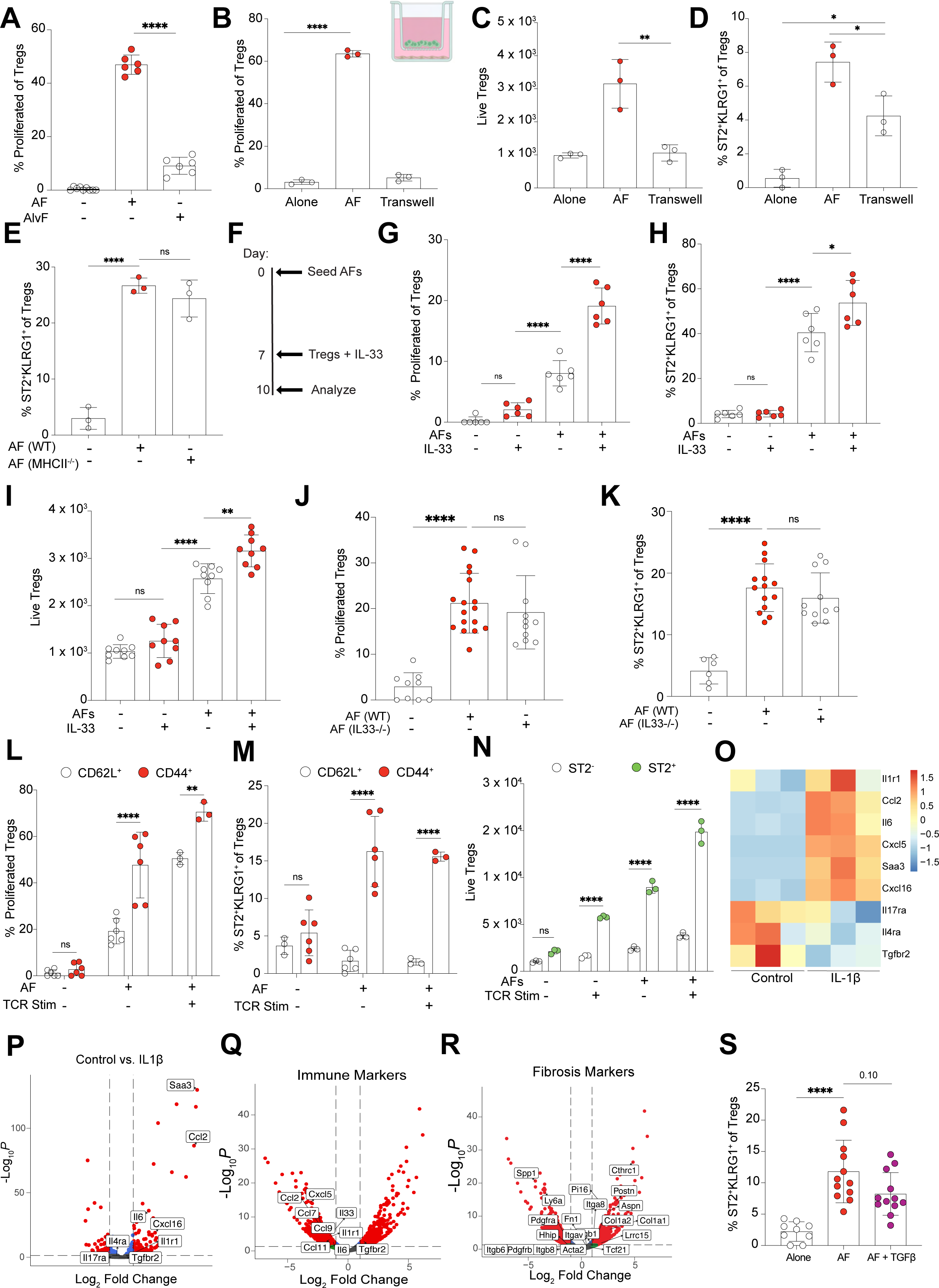
**A**, Quantitation comparing effects of AFs and AlvFs on Treg proliferation. **B-D**, Quantitation looking at effect of transwell cultures of AFs on Treg (**B**) proliferation, (**C**) viability, and (**D**) ST2^+^KLRG1^+^ expression. **E**, Quantitation looking at the effect of MHC II^-/-^ AFs on Treg ST2^+^KLRG1^+^ expression. **F-I**, (**F**) graphical illustration showing timeline of IL-33 and AF treatment and their effects on Treg (**G**) proliferation, (**H**) ST2^+^KLRG1^+^ expression, and (**I**) viability. **J-K**, Quantitation of effect of IL-33^-/-^ AFs on Treg (**J**) proliferation and (**K**) ST2^+^ KLRG1^+^ expression. **L-M**, Quantitation comparing AF support on effector (CD44^+^) and naïve (CD62L^+^) Treg (**L**) proliferation and (**M**) ST2^+^KLRG1^+^ expression. **N**, Quantitation comparing the effects of a combination of AFs and stim on ST2^+^ and ST2^-^ Treg viability. **O**, Heatmap of control AFs or AFs cultured with IL-1β illustrating markers associated with inflammatory fibroblasts. **P**, Volcano plot comparing DEGs of control and IL-1β-treated AFs (10ng/mL). **Q-R**, Volcano plots comparing DEGs of control and TGFβ-treated AFs (10ng/mL), looking at (**Q**) immune and (**R**) fibrosis markers. **S**, Quantitation comparing effects of AFs and TGFβ-treated AFs on Treg ST2^+^KLRG1^+^ expression. Graphs show mean±s.d. ns = not significant, *p<0.05, **p<0.01, ***p<0.001, ****p<0.0001; one-way ANOVA, Tukey post-test (**A-E, G-K, S**); two-way ANOVA, Sidak’s post-test (**L-N**).

**Figure S6:**
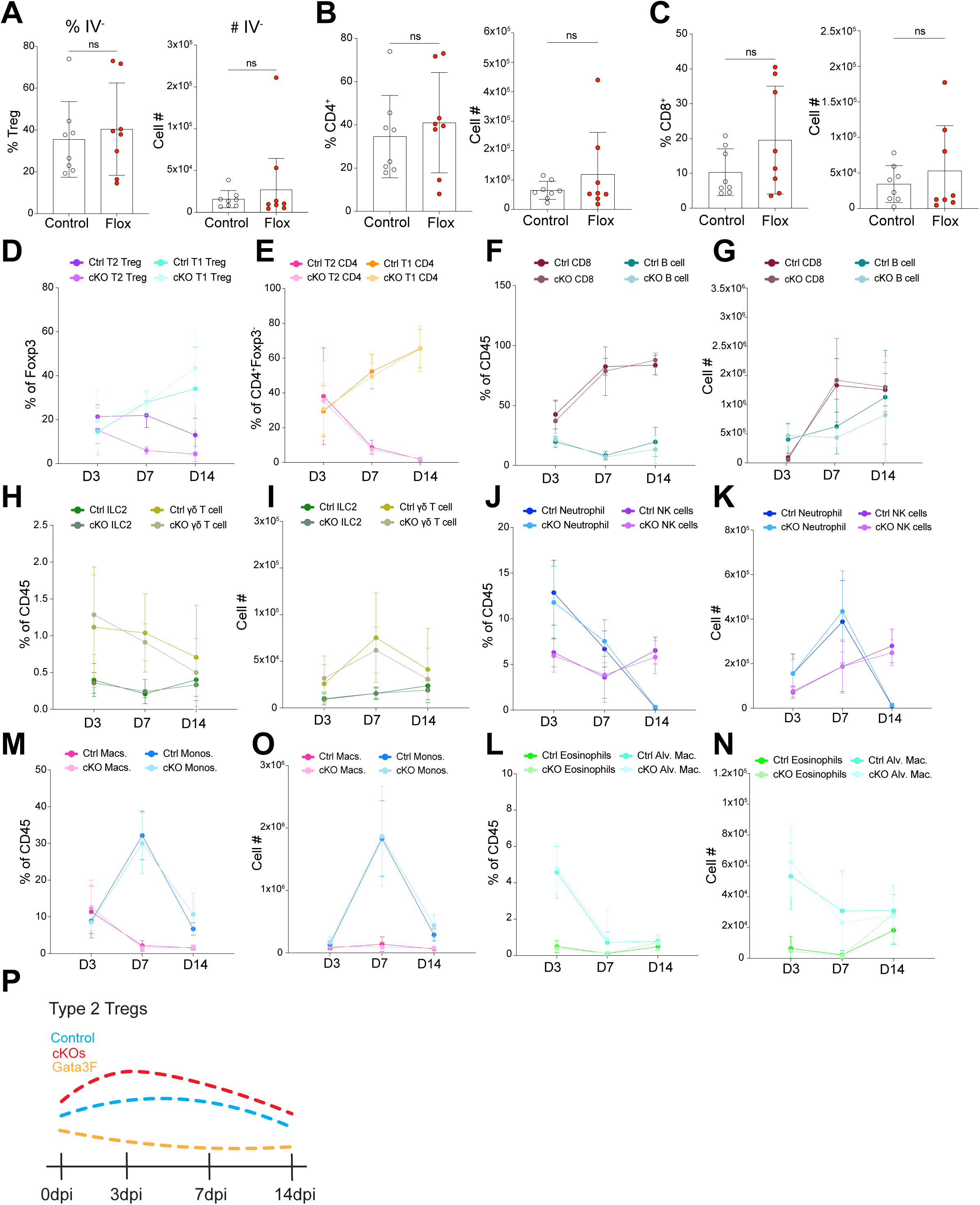
**A-C**, Quantitation showing frequency (left panels) and numbers (right panels) of tissue resident (i.v.^-^) (**A**) Tregs, (**B**) CD4^+^ T cells, and (**C**) CD8^+^ T cells. **D-E**, Quantitation showing frequencies of (**D**) T2L and T1L Tregs and (**E**) T2L and T1L CD4^+^ T cells. **F-N**, Time course showing frequency and numbers of various immune cell populations between control and flox mice at 3, 7, and 14 dpi. (**F, H, J, M, L**) Frequency of total CD45^+^ cells of (**F**) CD8 T and B cells, (**H**) ILC2s and ψ8 T cells, (**J**) Neutrophils and NK cells, (**M**) monocytes and macrophages, and (**L**) Eosinophils and alveolar macrophages. (**G, I, K, O, N**) Numbers of (**G**) CD8 T and B cells, (**I**) ILC2s and ψ8 T cells, (**K**) Neutrophils and NK cells, (**O**) monocytes and macrophages, and (**N**) Eosinophils and alveolar macrophages. **P**, Graphical illustration showing Type 2-like Treg dynamics in control, TGFbfr2 cKO, and T2 Treg deleter models. Graphs show mean±s.d. ns = no significance; Student’s T-test (**A-C**).

